# A comprehensive genetic catalog of human double-strand break repair

**DOI:** 10.1101/2024.08.03.606369

**Authors:** Ernesto López de Alba, Israel Salguero, Daniel Giménez-Llorente, Ángel Fernández-Sanromán, Ester Casajús-Pelegay, José Terrón-Bautista, Jonathan Barroso-González, Juan A. Bernal, Geoff Macintyre, Rafael Fernández-Leiro, Ana Losada, Felipe Cortés-Ledesma

## Abstract

The analysis of DNA sequence outcomes provides molecular insights into double-strand break (DSB) repair mechanisms. By employing parallel in-pool profiling of Cas9-induced indels within a genome-wide knockout library, we present a comprehensive catalog detailing how virtually every human gene influences the DSB repair process. This REPAIRome resource is validated through the identification of novel mechanisms, pathways and factors involved in DSB repair, including unexpected opposing roles for XLF and PAXX in DNA end processing, a molecular explanation for Cas9-induced multi-nucleotide insertions, the identification of HLTF as a DSB-repair factor, the involvement of the SAGA complex in microhomology-mediated end joining, and importantly, an indel mutational signature linked to VHL loss, renal carcinoma and hypoxia. Collectively, these results exemplify the potential of REPAIRome to drive future discoveries in DSB repair, CRISPR-Cas gene editing and the etiology of cancer mutational signatures.

## Introduction

DNA double-strand breaks (DSBs) are particularly dangerous lesions that disrupt the continuity of the DNA molecule. They represent important threats for genome integrity and are linked to severe pathologies such as cancer, immune and neurological disorders (*1–3*). Furthermore, DSB-inducing agents as well as inhibitors of DSB-repair factors and pathways are widely used in cancer therapy (4). More recently, the development of CRISPR-Cas methodologies for gene editing has further expanded the general interest in DSB repair mechanisms. For template-free gene editing, Cas9 cleavage is directed to the genomic locus of interest through the formation of a ribo-nucleoprotein complex (RNP) with a customizable guide RNA (gRNA) that drives base complementarity with the target sequence. This induces recurrent Cas9 cutting at the target site until mutagenic repair by non-homologous end joining (NHEJ) or microhomology-mediated end joining (MMEJ) results in insertions and/or deletions (indels) that prevent further sequence recognition and cleavage (5). Despite the uncontrolled editing outcomes, targeted indel induction remains a preferred method for generating gene knockouts due to its simplicity and high efficiency, compared to other CRISPR-based more precise technologies such as homology-directed repair and base or prime editing. A comprehensive understanding of how DSB repair mechanisms operate to give rise to specific mutational outcomes is therefore an area of extraordinary interest with profound implications for human health, including cancer biology and treatment, as well as in our efforts towards a full control over CRISPR-Cas gene-editing technologies.

Conversely, CRISPR-Cas systems provide an excellent tool to explore the molecular details of DSB repair mechanisms through the analysis of repair profiles in different contexts and genetic backgrounds (*6*). This, when combined with in-pool genetic screening, has fueled massive parallel interrogation of different aspects of Cas-induced DSB repair. Thus, it is now clear that the repair profile (i.e., the specific distribution of repair outcomes) is not random, but strongly determined by the local sequence context at the cut site, allowing for the development of accurate indel prediction algorithms that are routinely used in the design of specific gene-editing strategies (*7–10*). Moreover, the differential engagement of DSB repair pathways and factors, i.e., the processing of DNA ends by DNA polymerases or nucleases, together with the usage or not of base pairing to bridge DNA ends across the break in NHEJ and MMEJ, constitute the ultimate effectors responsible for each given repair event, and can be therefore modulated to specifically control gene editing outcomes (*6*). A recent genetic screening linking the specific knockdown of more than 400 known DSB repair-related factors to their effects on Cas-induced DSB repair outcomes has provided a remarkable proof-of-concept on how similar repair events have common genetic requirements, while factors belonging to the same pathway similarly affect repair patterns (*11*). This seminal study (Repair-seq) highlights the potential of massive parallel strategies to gain novel insights into pathways affecting DSB repair and CRISPR-Cas gene editing, but the use of a focused library limited the discovery of novel DSB-repair factors and mechanisms. Although reporter substrates have been used to increase the throughput capacity and allow interrogation of genome-wide perturbation libraries (*12*), this is done at the cost of losing nucleotide-level molecular resolution achieved by sequencing.

Here, we optimize in-pool DSB repair sequence profiling, significantly increasing its throughput capacity to tens of thousands genetic conditions that are sufficient for whole-genome coverage. With this genome-wide screen, we obtain a comprehensive resource of how each of more than 18-thousand human gene affects repair of Cas9-induced DSBs (REPAIRome). This “genetic catalog of DSB repair” can be consulted for any gene of interest, as well as for the discovery of potentially novel relationships and pathways, with the help of a publicly available and browsable webtool (will be accessible upon publication of the work). Furthermore, initial examination of the vast amount of data generated has led us to the identification of new DSB repair factors and the discovery unexpected functions and genetic interactions of known repair proteins. It has also provided new insights regarding Cas9-mediated cleavage with implications for gene-editing outcomes, and even uncovered the molecular etiology of an orphan indel signature found in cancer. These results highlight the potential and usefulness of the rich resource generated to foster discoveries in DSB repair and CRISPR-Cas gene editing mechanisms.

## Results

### Genome-wide genetic profiling of DSB repair outcomes

Following a similar pipeline to previous DSB repair-profiling studies focused of a few hundred DNA repair and related factors (*11*), we based our strategy on a lentiviral sgRNA genetic perturbation library singly integrated within the genome of the cell, and then targeting a CRISPR-Cas9-induced DSB to the common backbone of the construct, in the vicinity of the sgRNA-expressing cassette (figure 1A). Thus, the repair event occurring at the library target site and the specific genetic condition are molecularly linked, so they can be read and individually matched by subsequent targeted amplification and Illumina sequencing. With this information, indel repair profiles for each of the genetic perturbations in the library can be computed and analyzed (figure 1A). To expand the throughput capacity to the genome-wide level, the choice of an optimal genetic perturbation system and library is essential to maximize the output and keep library representation (number integrations per sgRNA) and experimental depth (total number of repair events per sgRNA) within what is experimentally feasible. We decided to use the TKOv3 library, which includes lentiviral expression vectors for 70948 sgRNAs targeting 18052 human genes, and 142 additional non-targeting controls, in p53-knockout *hTERT* immortalized RPE1 cells overexpressing the SpCas9 protein (RPE1-Cas9 *TP53^-/-^*). Importantly, the *TP53* mutation avoids losing DSB repair factors due to decreased cellular fitness in this non-transformed cell line but does not affect CRISPR-Cas-induced indel distribution (figure S1A). Constitutive Cas9 overexpression results in high knock-out efficiency, above 70% for most sgRNAs in the library, that produce stronger and more readily detectable phenotypes than depletion, making it an ideal setting for in-pool CRISPR genetic screening to dissect the cellular response to DNA damaging agents (*13*, *14*).

**Fig. 1.**
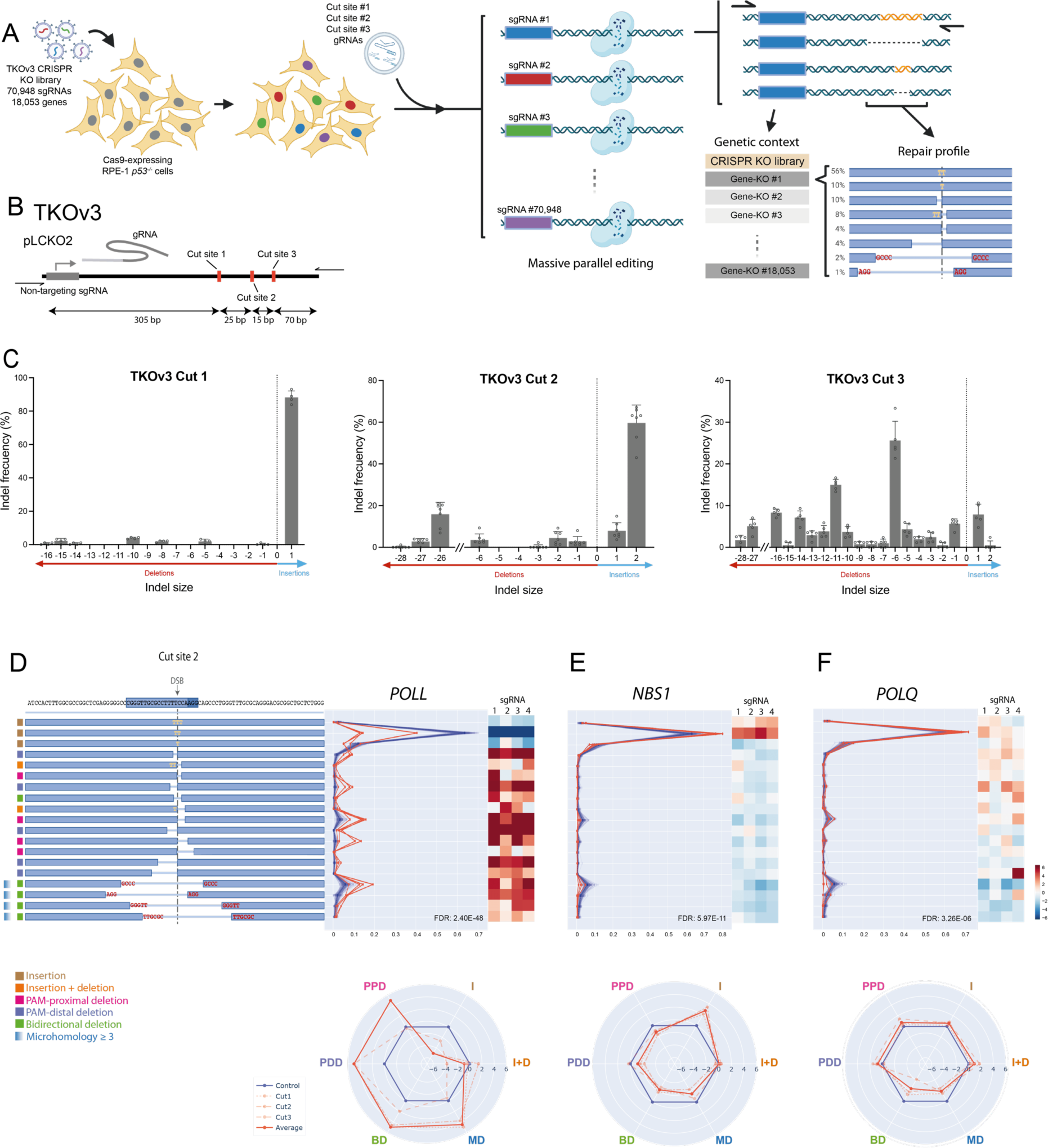
REPAIRome screen design and setup. **(A)** Experimental set up of the screen. Cas9-experssing *p53^-/-^* hTERT RPE-1 cells are infected with the TKOv3 lentiviral library. After puromycin selection, infected cells are kept growing for 12 days and then transfected with a gRNA targeting a region of the lentiviral backbone in the proximity of the library sgRNA sequence. Cells are allowed to repair the DSB for 72 hours before genomic DNA purification and amplification of the region containing the library sgRNA sequence and the associated repair outcome. Illumina libraries are then prepared from this amplicon and pair-end sequencing is carried out to link each gene knockout to the DSB repair outcome. **(B)** Schematic representation of the TKOv3 sgRNA expression cassette. Cut sites used in the screen and location of primers used to generate the Illumina library are indicated. **(C)** Repair patterns of DSBs generated after transfection of RPE1-Cas9 *TP53^-/-^* TKOv3 cells with each of the three gRNAs used in the screen (see Methods). The relative abundance of each indel is normalized to the total percentage of edited sequences and then classified and ordered by size to facilitate the visualization. Negative indels refer to deletions (red arrow) and positive indels refer to insertions (blue arrow). Data show average of indel frequency, with error bars for ±SD (n= 4 for Cut site 1, n= 7 for Cut site 2, n=5 for Cut site 3). **(D, E and F)** Top left. Diagram showing the most frequent indels analysed in the screen for cut site 2. Protospacer and PAM sequences are highlighted in blue on the top. The expected cut site for each sgRNA is indicated by a vertical dashed line. Inserted nucleotides are shown in yellow and microhomology regions flanking a deletion are shown in red. The category of each indel is indicated by a coloured square on the left side of the diagram. Top centre. Frequency of each indel, relative to the total frequency of edited events, for the control (blue lines) and the indicated gene sgRNAs (red). Top right. Heatmap showing the Log2 fold change for each indel of the indicated sgRNA relative to the average of control sgRNAs. Bottom. Radar plots for the change of each indel category in the indicated genetic backgrounds. Values show normalized frequency change of indicated sgRNAs relative to controls (blue line). The continuous red line shows average change of the three cut sites combined, while the dashed lines show changes for each independent cut site.

Besides obtaining a rich pooled population of individual knock-out cells, another essential step is to achieve efficient Cas9 cleavage/editing in the subsequent step of target DSB induction. Editing efficiencies above 80% are typically obtained in RPE1-Cas9 *TP53^-/-^* cell lines by a simple transfection of tracrRNA-crRNA complexes (gRNA from now on). Importantly, the constitutive expression of a non-targeting sgRNA from the TKOv3 library construct (TKOv3-nt) still allowed high on-target editing efficiencies following gRNA transfection (figure S1B), suggesting that Cas9 levels are not limiting. The RPE1-Cas9 *TP53^-/-^* TKOv3 system is therefore compatible with induction of target cleavage by gRNA transfection. We used three gRNAs directed to a region downstream of the expression cassette in the library (figure 1B), each with a different predicted pattern of low (Cut 1), mid (Cut 2) and high (Cut 3) variability of repair outcomes (as determined by the in-Delphi predictor (*8*). A high editing efficiency (between 75 and 95 %) (figure S1C) and the predicted repair patterns (Cut 1: strongly predominant +1 insertion; Cut 2: predominant +2 and +1 insertions together with specific deletions; Cut 3: range of different insertions and deletions) were obtained (figure 1C). We reasoned that the combined outcomes of these three cut-sites would provide a representative picture of the contribution of the main end-joining DSB-repair pathways.

Finally, we performed a pilot experiment to determine the library representation and experimental depth required to obtain reproducible indel patterns. We used a barcode lentiviral library (Clone-tracker library, Cellecta) conferring no genetic perturbation, and obtained a pooled population of around one thousand individual clones harboring a unique integration. Two editing experiments were performed, each with a different gRNA directed to a region adjacent to the library barcode (figure S1D). Barcode and target sites were amplified and massively sequenced. With these data, we performed computational analyses to determine the variability (error) in the pattern of indels following 100 virtual random iterations for different combinations of a fixed number of barcodes (representation) and total reads (depth) (figure S1E). We observed that depth was the strongest determinant, continuously decreasing experimental variability up to the 2,000 total events tested. In contrast, beyond 50 independent barcodes, there was little contribution of increasing representation, suggesting that this number may be sufficient to buffer the positional variability of random lentiviral integration. From these results, we concluded that a 500x library representation, which is necessary to maintain library variability, and above 1000x total depth were sufficient to obtain robust indel patterns; numbers that are compatible with the around 80,000 constructs of the TKOv3 library.

### REPAIRome: a genetic catalog of human CRISPR-Cas-induced DSB repair

We next conducted the genome-wide screen with the TKOv3 library in optimized conditions described above, with two experimental replicates for each of the three cut-sites tested. Once all the sequencing results were obtained, filtered and matched, we computed, for each sgRNA in the library, the frequency of the 65 most abundant outcomes (62 indels and 3 non-edited events) and the corresponding normalized difference from non-targeting controls (see methods), obtaining the distribution of DSB-repair events for each genetic perturbation and how this changed from unperturbed conditions. In order to appreciate changes in the editing patterns, irrespectively of potential effects on overall editing efficiencies, the distribution of indels and the respective difference from controls were also calculated without considering non-edited events. Aggregated frequencies were also computed for total editing efficiency, insertions and deletions, as well as for specific sub-categories of events such as protospacer adjacent motif (PAM)-proximal deletions, PAM-distal deletions, bidirectional deletions, deletions with microhomology and combined insertion-deletions (figure S1F).

We then used the median values of the corresponding individual sgRNAs to calculate event and indel distributions, as well as differences to controls, for each of the genes. Then, to obtain a single value representing the overall difference in repair outcome, we calculated the Euclidian distance to controls of the entire indel distribution (“distance” from now on), which ranged from 0 to more than 50 (table S1). As an indication of the robustness of the observed differences, we tested, for each cut site and replicate, whether the values of the set of sgRNAs targeting each gene were significantly different to those of non-targeting sgRNA controls (false discovery rate; FDR, see methods), as well as the Pearson Correlation Coefficient (PCC) between the replicates. Finally, to explore potential relationships, we calculated the PCC of indel frequencies of genes with each other. To facilitate the visualization and analysis of the results, we have generated a dedicated webtool (will be accessible upon publication of the work) (see figure 1D-F for examples).

### Global analysis of REPAIRome results

To validate the utility of the resource to obtain novel insights into Cas9-induced DSB repair, we focused on the set of 168 genes with the strongest impact on the pattern of indel distribution (table S2). These were identified as those with a distance > 5 and in which, at least for one of the cut sites, the indel distribution was significantly different to controls in the two replicates (FDR < 0.01), which also showed correlation between replicates (PCC > 0.3) (see figure S2A for an example of Cut 2). STRING protein-protein interaction enrichment analysis suggested strong functional connections between the selected genes (p < 1e-16) (figure S2B and S2C), supported by significantly enriched gene-ontology terms “DSB repair” (GO:0006281; n= 22; FDR= 2.1e-13), “DNA repair” (GO:0006281; n=27; FDR= 1.3e-11) and “DSB repair by NHEJ” (GO:0006303; n=12; FDR= 2.2e-11). 135 of the selected genes were not associated with these GO terms (figure S2B, white circles), but were also functionally inter-related (p = 4.62e-07), suggesting potentially unknown factors and pathways.

To better understand the relationships between the selected candidate genes, we calculated the correlation among the 65 events and the 168 genes, and then performed unsupervised hierarchical clustering (figure 2A). Events of the same type (as categorized in figure S1F) had the tendency to cluster together, indicating that they had common genetic requirements, as previously demonstrated (*11*). For further analysis and visualization of the selected genes, we performed UMAP dimensionality reduction, in which genes with similar distribution of events cluster in the vicinity of each other, while different variables of interest can be additionally plotted on the generated maps (figure 2B-E). A representation of distance (figure 2B and table S2) highlighted a cluster of genes containing well-established NHEJ factors, such as LIG4, XRCC4, XLF and POLL, together with DDX5 and ERCC6L2, which have been related to NHEJ recently (*14–16*), and several components of the neddylation machinery (CUL3, UBE2M, UBA3 and KCTD10), which is known to promote removal of the Ku complex from DNA ends (*17*) Surprisingly, this cluster also contained HLTF (distance = 19.7), a factor involved in the regulation of DNA damage tolerance pathways and replication-fork reversal that has not been previously linked to DSB repair (explored further below) (*18–22*). Finally, there were two additional genes showing a high (>10) distance that did not cluster with classic NHEJ factors: APTX, which is known to clean-up adenylated DNA ends and serve as a proof-reading mechanism for ligation events (*23–26*), and FBXL12, which promotes degradation of repair proteins such as FANCD2 and Ku80 (*27*, *28*). The molecular mechanism responsible for their effect on DSB-repair outcomes is an interesting avenue for future research.

**Fig. 2.**
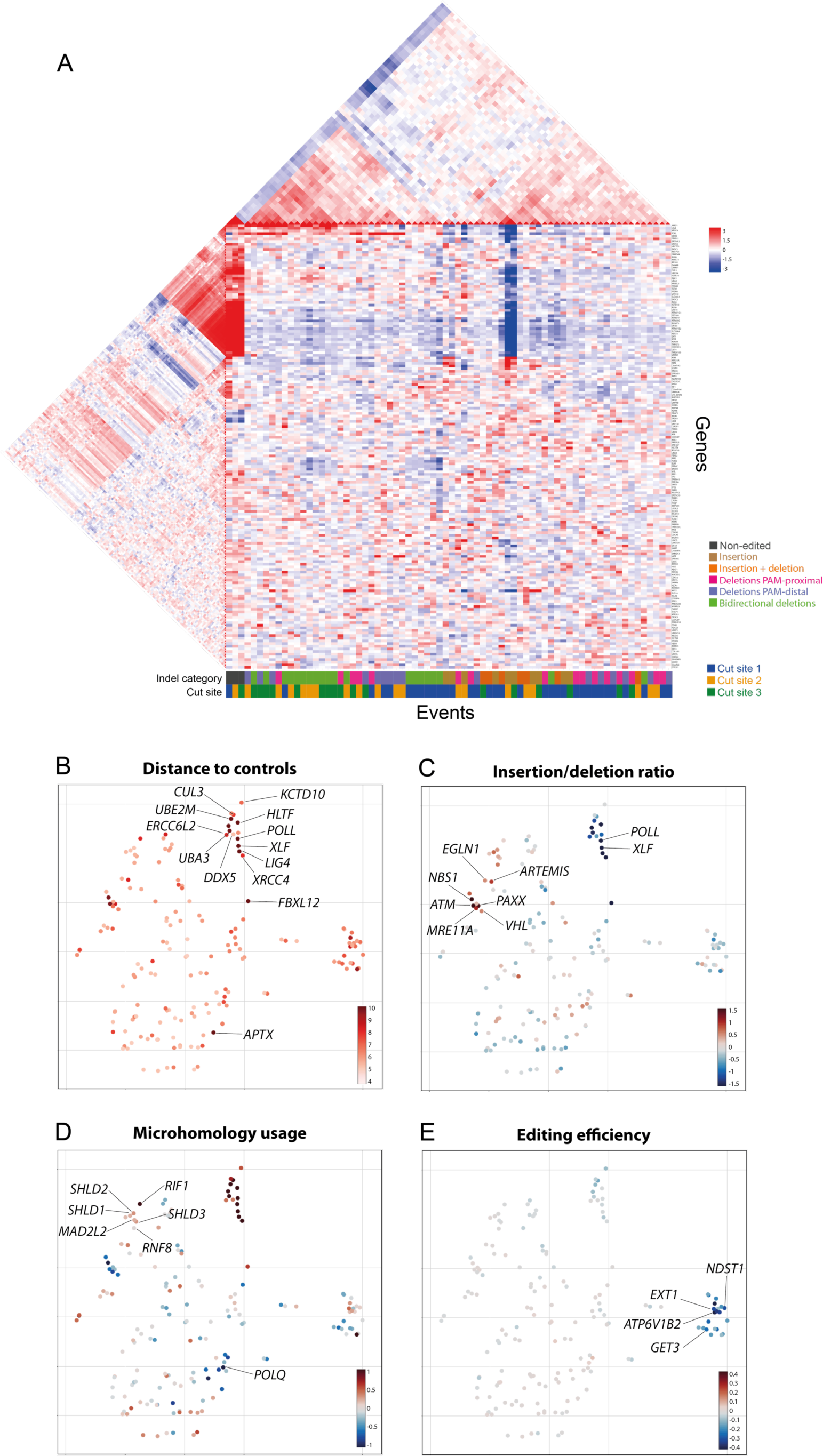
Global analysis of REPAIRome results. **(A)** Heatmap showing Log2 fold changes of all indels for the selected genes relative to control sgRNAs. Unsupervised clustering was performed for genes and indels. On the central part of the map rows correspond to median values of gRNAs for each gene and columns correspond to individual indels coming from the three cut sites of the screen. Triangular heatmaps show correlation between events (above) and between genes (left). Indel categories and corresponding cut site for each event are indicated by coloured squares on the bottom. **(B-E)** UMAP dimensional reduction of indels pattern alteration for selected genes. Analysis of additional parameters as distance to repair profiles of control sgRNAs (B), insertions vs deletions ratio (C), microhomology usage (D) and editing efficiency (E), are overlayed on the generated UMAPs.

Next, we explored the ratio between insertions and deletions (figure 2C), for which the NHEJ cluster displayed the lowest values, particularly the DNA polymerase *POLL* [log2(insertion/deletion) = −8.6] (see also figure 1D and table S2). This indicates that the insertions observed at these cut sites (fundamentally +1 and +2) occurred via POLL-mediated NHEJ, likely through the fill-in of staggered Cas9-induced DSBs, as suggested previously (*29*, *30*). Knock-out of main DSB repair-related nucleases, such *ARTEMIS* and members of the MRN complex *MRE11* and *NBS1* (see also figure 1E), strongly increased the insertion/deletion ratio, consistent with deletions requiring, as expected, nucleolytic trimming of DNA ends. *ATM* also clustered within this region, in agreement with its main repair function being related to the regulation of end processing (*31*). *PAXX*, a paralog of *XLF*, with whom it is thought to redundantly favor ligation during NHEJ (*32–37*), unexpectedly clustered with the deletion-promoting genes, instead of with the insertion-promoting NHEJ cluster (explored further below). Interestingly, this cluster also contained *VHL* and *EGLN1*, regulators of the hypoxia response with relevant implications in cancer, but with no reported DSB-repair function (*38*).

*POLQ*, a key factor in MMEJ, was also among the 168 selected gens. As expected, it strongly limited the usage of microhomology but had only minor effects on other events (figure 2D; table S2). This was different from the profile obtained for knockout of end-processing factors such as *NBS1*, that reduced, not only microhomology-dependent deletions, but all types of deletions (compare figures 1E and 1F). In addition, the entire Shieldin complex (*SHLD1*, *SHLD2*, *SHLD3* and *MAD2L2*), which has end-protection functions during DSB repair (*39–44*), was found, together with *RNF8* and *RIF1*, in a separate cluster of genes whose loss caused increased microhomology usage, while increasing the insertion/deletion ratio (figure 2C and D). Altogether, these results validate the capacity of the REPAIRome resource to identify DSB repair factors and to distinguish between the different pathways and processes involved.

Finally, we noted a separate cluster of knockouts that were characterized by causing a marked decrease in the overall editing efficiency, i.e., increased frequency of non-edited events (figure 2E). A STRING analysis of these genes showed highly significant functional relationships (p < 1e-16) related to the cellular endomembrane system (figure S2D). This finding may be relevant for CRISPR-Cas9 biotechnological and clinical applications, and has also allowed us to explore how editing efficiency affects indel distribution (see below). To further support the robustness and power of the REPAIRome screen, the following sections include validation and functional characterization of some of the most interesting observations described above.

### XLF and PAXX opposingly control end processing by regulating NHEJ synaptic structures

As an example of potential new functions of known DSB repair factors, we focused on the differential repair profiles caused by loss of XLF and PAXX paralogs, which conferred a clear bias towards deletions and insertions, respectively (figure 2C, 3A). We first validated these initial observations and ruled out a contribution of the *TP53^-/-^* background using pooled populations of p53-proficient RPE1-Cas9 TKOv3-nt cells transfected with *de novo*-designed sgRNAs targeting *XLF* and *PAXX* (table S3 for knock-out levels), and assessing repair at Cut 2 (figure S3A). We then analyzed two independent mutant clones for each of these genes (figure S3B, table S5), which showed a consistent and more pronounced effect than the polyclonal populations, with virtually no insertions and increased deletions in the *XLF^-/-^* mutants, and the opposite for *PAXX^-/-^* (figure 3B). This differential behavior was additionally validated at two different cut sites within the *AAVS1* endogenous locus (figure S3C and D). These results demonstrate unexpected opposing functions of the two paralogs in DSB repair, which likely result from XLF and PAXX favoring fill-in and nucleolytic trimming of DNA ends, respectively. Indeed, indel patterns in the absence of POLL, the polymerase likely responsible for the fill-in reaction, displayed a dramatic reduction in insertions, similar to the *XLF^-/-^* mutant, both in a polyclonal population (figure S3A; knock-out levels in table S3) and in two *POLL^-/-^* clones (figure 3B; validation in figure S3B, table S5). The distribution of deletions, however, was clearly different between the two mutants: preferentially long, microhomology-mediated in *XLF^-/-^*, and short, microhomology-independent in *POLL^-/-^*; signatures that were already noticeable in the results of the screen (POLL in figure 1D and XLF in figure 3A). These results suggest that XLF not only promotes fill-in during repair by NHEJ, but also prevents nucleolytic trimming. Finally, the *XLF^-/-^* mutation was clearly epistatic over *PAXX^-/-^*, with the phenotype of the double mutants being nearly identical to that of the *XLF^-/-^*singles (figure 3B), suggesting that PAXX mainly operates by counteracting XLF function.

**Fig. 3.**
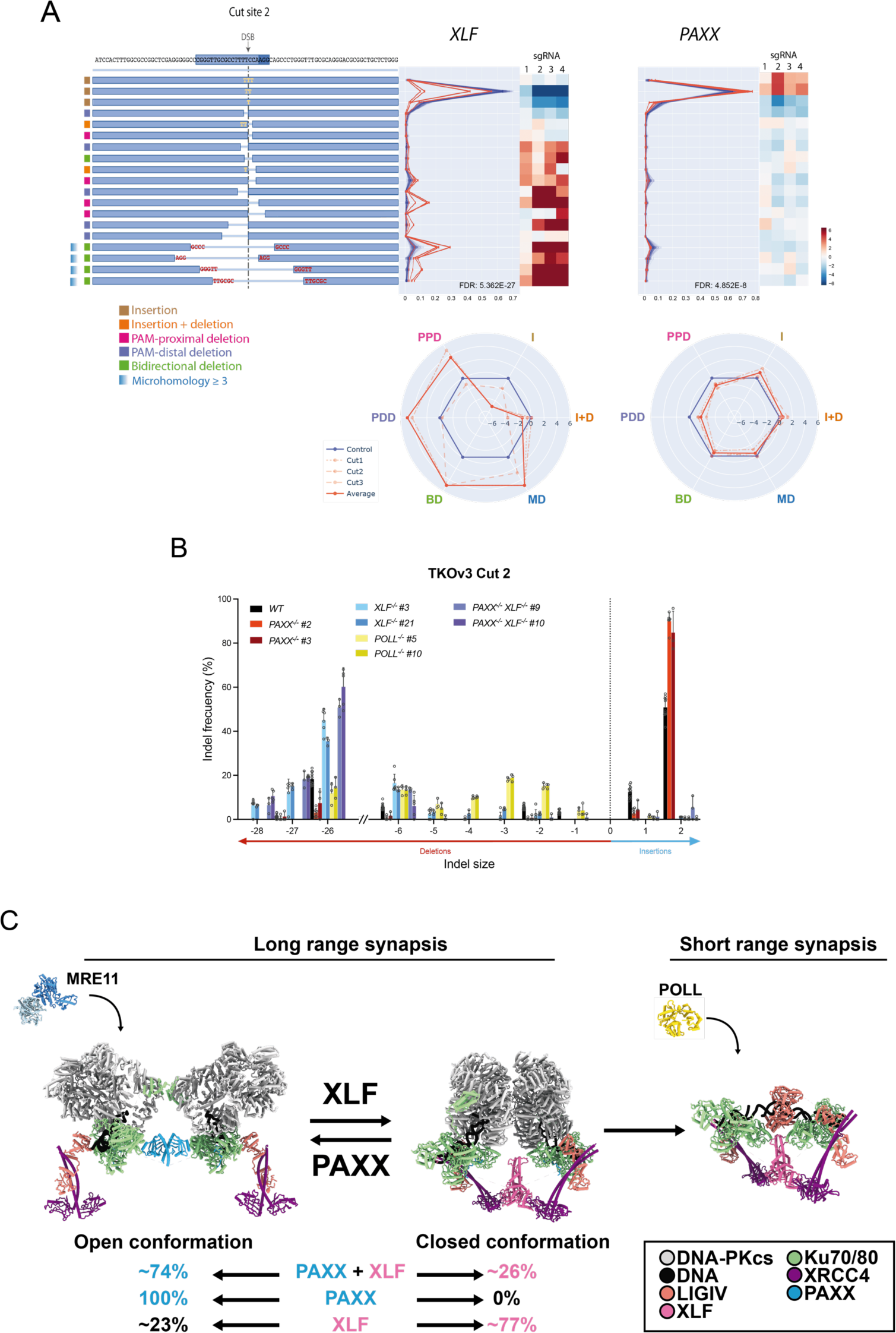
XLF and PAXX opposingly control end processing by regulating NHEJ synaptic structures. **(A)** Summary of results obtained in the screen for XLF and PAXX, as in (figure 1D, E and F). Only TKOv3 Cut 2 is shown. **(B)** Impact of XLF, PAXX and POLL absence on the repair profile at TKOv3 Cut site 2 in the indicated RPE1-*Cas9* TKOv3-NT clones. Experiments are carried out as in figure 1C. Data show average values normalized to total editing efficiency. n = 6, error bars show ±SD. **(C)** Model for PAXX/XLF opposing effect on the equilibrium between open and closed conformations of NHEJ synapses. Particle distribution information from published cryo-EM studies (*36*, *37*, *46–48*) was used to evaluate the effect of PAXX and XLF on the equilibrium and is shown here as a percentage of the total long-range particles found in those datasets (see Methods for a description of data analysis and selection). Representative structures for the long-range open (8BH3) and closed (8BHV) conformations, short-range complex (7LSY), MRE11 (1XSP) and POLL (8BAH) are shown to illustrate the model.

In recent structural models of NHEJ, three synaptic configurations have been identified (figure 3C) (*36*, *37*). First, there is an “open” long-range structure mediated by *in trans* Ku80-DNA-PKcs interactions at either side of the DSB. Second, in the presence of XRCC4, LIG4 and XLF, the long-range complex can be also organized as a “closed” configuration that is mediated by XLF and DNA-PKcs dimerization across the break, in equilibrium with the open long-rage arrangement. Last, the closed long-range structure can transition into a ligation-compatible short-range complex mediated by XLF, and not DNA-PKcs. Notably, PAXX was recently found in open long-range synapses in the absence of XLF, and both in open and closed complexes when XLF was present, which was interpreted as proof for their reported functional redundancy (*36*, *37*). The open long-range structures appear to favor nucleolytic accessibility, while the formation of short-range complexes is required for end fill-in (*45*). We therefore reasoned that XLF and PAXX could affect the repair outcome by regulating the transitions between synaptic structures. To test this hypothesis, we used published Cryo-EM data to determine how XLF and PAXX affect the distribution of particles in open and closed synaptic conformations (figure 3C, table S4) (*36*, *37*, *46–48*). In agreement with the phenotypes and genetic relationships observed, the presence of PAXX strongly shifted the equilibrium of XLF-containing complexes from closed to open structures in these cryo-EM datasets (table S4). Altogether, these results suggest a mechanism by which PAXX is redundant with XLF to promote the formation of open long-range complexes, but restrains XLF-mediated formation of the closed long-range configuration and subsequent transition into short-range synapsis, thus favoring nucleolytic processing and inhibiting fill-in of DNA ends.

### Target re-cleavage explains multi-nucleotide insertions

As mentioned above, we found a cluster of genes generally related to cellular endomembrane systems (figure S2D) that, when deleted, strongly decreased overall editing efficiency (figure 2E and S4A). This observation was validated in pooled knock-out p53-proficient populations of three genes, *GET3*, *NDST1* and *ATP6V1B2* (figure S4C; table S3 for knock-out levels), each representing one of the main pathways involved. In the case of *GET3* we also tested two independent clones (validated in figure S4B, table S5), which displayed a similar reduction in the percentage of edited events (figure 4A). We reasoned that, for these repair-unrelated cytoplasmic factors, a reduction in editing efficiency could be caused by a reduction in Cas9 cutting efficiency. Indeed, both *GET3^-/-^*clones showed reduced on-target cleavage, as determined by qPCR across Cut 2 of the library (figure 4B) and a decrease in the induction of 53BP1 foci (a well-established marker of DSBs) (*49*) when transfected with a gRNA targeting over 100 sites in the genome (figure 4C and S4D) (*50*). This phenotype is not caused by a reduction in Cas9 cellular levels (figure S4E), but may be a consequence of changes in the uptake, intracellular distribution and/or stability of the sgRNA. Although far from our direct interest in DSB-repair pathways, this finding is relevant for the biotechnological and medical applications of CRISPR technologies.

**Fig. 4.**
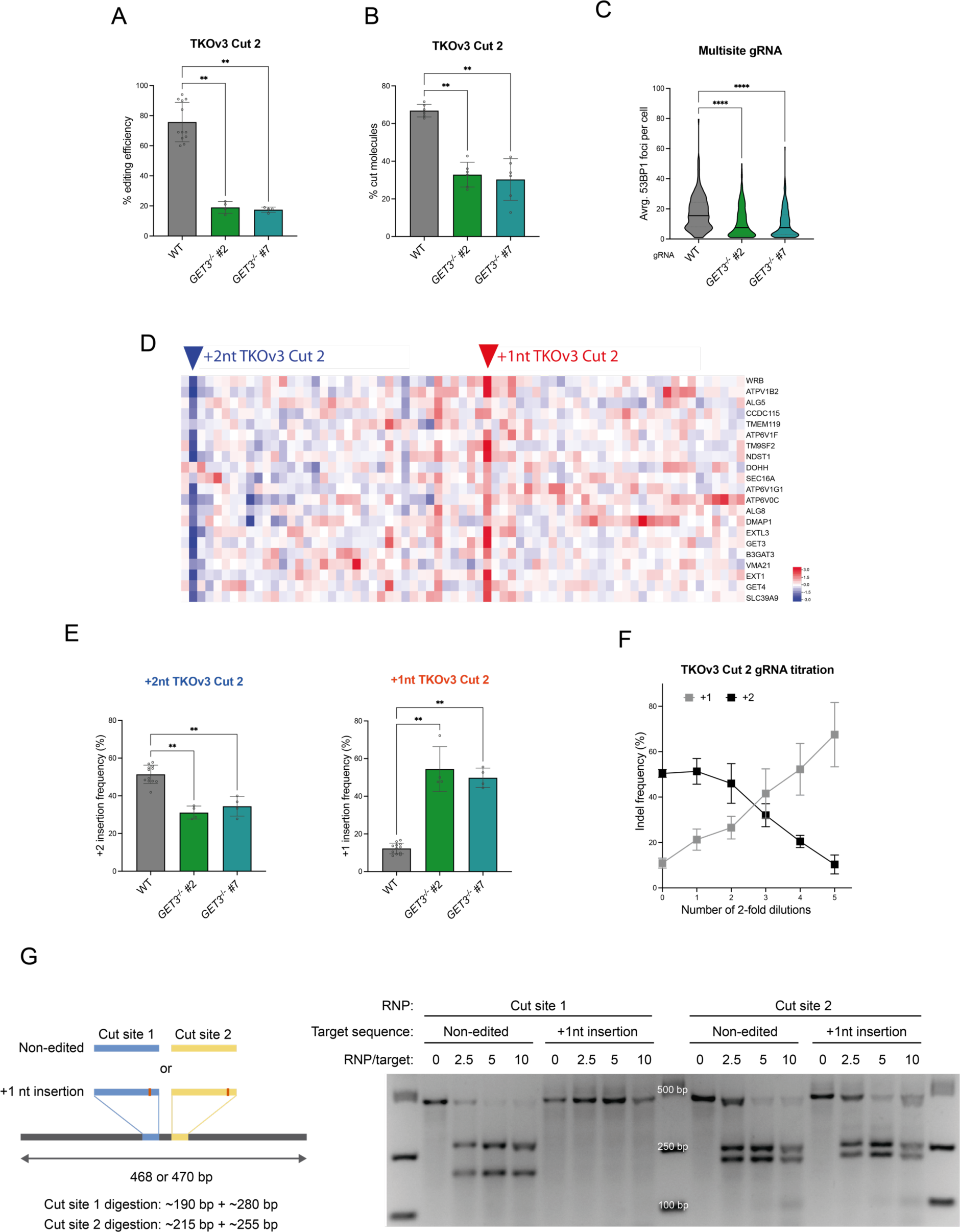
Target re-cleavage explains multi-nucleotide insertions. (A) Effect of *GET3* on overall editing efficiency at TKOv3 Cut 2 in RPE1-*Cas9* TKOv3-NT cells. Experiments carried out as in figure 1C. Two *GET3^-/-^* clones are tested. n = 12 for WT and 4 for *GET3^-/-^* clones. Error bars show ±SD. Statistical analysis was performed using One-way ANOVA. (B) Cas9 cutting efficiency at TKOv3 Cut site 2 estimated by qPCR. Primers annealing at both sides of the cut site are used to determine the percentage of cut molecules 6h after gRNA transfection in the indicated cell lines. Amplification levels were normalized to the endogenous gene GREB1. n = 6, error bars show ±SD. Statistical analysis was performed using One-way ANOVA. (C) 53BP1 foci number per cell 12h after transfection with a multi-target gRNA (1.5 nM) in the indicated cell lines. Data shows foci count of two independent replicates merged. Statistical analysis was performed using One-way ANOVA. (D) Heatmap showing indel frequency variation (Log2 fold change) in the three cut sites of the screen for the reduced editing efficiency gene cluster (figure 2E). Two insertion events in the TKOv3 Cut site 2 that are consistently reduced or increased across most knockouts in this group are marked with a blue and a red triangle, respectively. (E) Frequency of +2 (left) and +1 (right) nucleotide insertions, relative to the overall editing efficiency, after transfection with the TKOv3 Cut site 2 gRNA in RPE1-*Cas9* TKOv3 *GET*3^-/-^ clones compared to wild type cells. Experiments carried out as in figure 1C. n = 12 for WT and 4 for *GET3^-/-^* clones. Error bars show ±SD. Statistical analysis was performed using One-way ANOVA. (F) Variation in the frequency of +1 and +2 nucleotide insertions produced after transfection with decreasing concentrations (2-fold serial dilutions from our standard gRNA final concentration, 12 nM) of gRNA targeting TKOv3 Cut site 2 in RPE1-*Cas9* TKOv3-NT Cells. n = 3, error bars show ±SD. (G) In-vitro Cas9 DNA cleavage assay showing differential gRNA mismatch tolerance. Schematic representation of the two constructs containing Cut site 1 and 2 sequence either unedited or including the most common +1 nucleotide insertion outcome observed in the screen data (left) (see Methods). Agarose gel showing digestion of the constructs with increasing concentrations of Cas9 ribonucleoparticle bearing gRNA targeting either Cut site 1 or 2 (right).

We then wondered whether the reduced editing efficiency of these mutants was also accompanied by changes in indel distribution, which would explain their selection under the criteria applied. Indeed, the results of the screen for these genes consistently showed decreased +2 and increased +1 events, specifically in Cut 2 (figure 4D), a result that was confirmed in pooled knockout populations of representative candidates (figure S4F) and in *GET3^-/-^* clones (figure 4E). Moreover, in wild-type cells, we also observed a clear and opposite dose-response effect on +2 and +1 insertions when the amount of Cut 2 gRNA was reduced (figure 4F), indicative of a strong link between these two indels and cutting/editing efficiency. A simple explanation for this could be that +2 events occur by two consecutive rounds of 1-nucleotide insertion after target re-cleavage. This would require the sequence of the +1 insertion to be also recognized and cleaved by the target Cas9 RNP. To test this, we generated DNA fragments containing Cut 2 (predominant +2) and Cut 1 (predominant +1) target sites, together with their +1 modified versions, and checked how the Cut 2- and Cut 1-specific Cas9 RNPs cleaved each fragment *in vitro* (figure 4G). As expected, both RNPs recognized and cleaved their unmodified target sites with similar high efficiency. However, the respective +1 versions were only efficiently cleaved in the case of Cut 2. We therefore conclude that at least some +2 insertions occur by a double +1 event, and that this is specific to particular target sequences in which +1 events can still be recognized and cleaved by the Cas9 RNP. This changes the view of multi-nucleotide insertions occurring by an alleged flexibility in the staggering of Cas9 cuts (*30*, *51*, *52*) and should be considered to improve current indel prediction algorithms.

### HLTF removes post-cleavage Cas9 RNP from DNA ends

*HLTF* was ranked third in overall distance to controls, just below *POLL* and *XLF*, and was thus the DSB repair-unrelated factor most strongly affecting repair outcome. The general effect was a decrease in insertions and an increase in deletions, but it was somewhat variable between the three cut sites, displaying the strongest effect in Cut 2 (figure 5A and S5A), which we confirmed in *HLTF* pooled (figure S5B and table S3 for knock-out levels) and three clonal (figure 5*B*; knock-out confirmation in figure S5C, table S5) knock-out populations. Consistent with this, the effect of decreased insertions and increased deletions was observed to a variable extent in four cut sites tested at the *AAVS1* locus in a *HLTF^-/-^* clonal population (figure S5D). HLTF is a multimodal protein, with RING, ATPase and HIRAN domains, that play distinct roles during replicative stress and DNA damage tolerance (*18*). The RING domain is responsible for PCNA poly-ubiquitination to prevent translesion synthesis and favor template switching mechanisms (*19*, *53*, *54*), while HIRAN-mediated binding to 3’-hydroxyl ends and ATPase-dependent helicase activities cooperate to promote replication fork reversal (scheme in figure S5E) (*20*, *21*, *55*, *56*). The observed changes in the indel profiles, however, were unrelated to replication, as they were also observed in serum-deprivation arrested cells, and also upon *PCNA* in-pool deletion (figure 5C), suggesting that they were also independent of PCNA ubiquitination. Then, to determine the contribution of the different domains of HLTF to the DSB repair phenotype, we complemented *HTLF^-/-^* cells (confirmation in figure S5C) with different versions of *HLTF* carrying already characterized mutations in the HIRAN (*HLTF-R71E*), ATPase (*HLTF-D557A,E558A*) and RING (*HLTF-C760S*) regions (indicated in figure S5E) (*22*) and checked their effect on the repair profile obtained at *AVVS1* Cut site 4. When compared to the empty-vector control, repair was reverted towards increased insertions upon expression of wild-type and the RING *HLTF-C760S* mutant, but not the HIRAN *HLTF-R71E* and ATPase *HLTF-DEAA* mutants (figure 5D). This demonstrates that HLTF function in DSB repair is not related to its ubiquitin-ligase activity, but instead depends on 3’-end binding and helicase activities, which are essential for fork reversal.

**Fig. 5.**
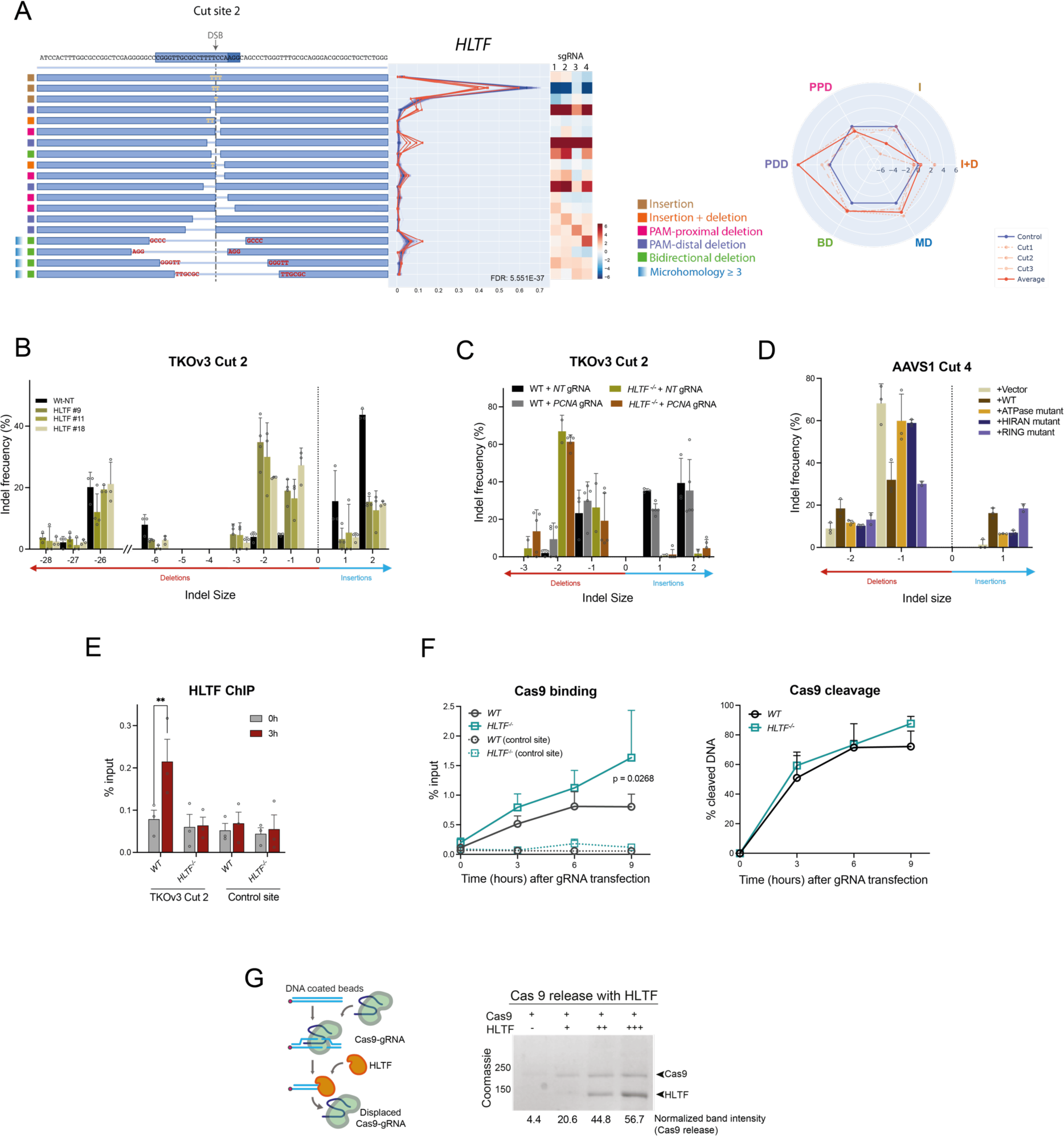
HLTF removes post-cleavage Cas9 RNP from DNA ends. **(A)** Summary of results obtained in the screen for HLTF, as in figure 1D. Only data for TKOv3 Cut site 2 is displayed. **(B)** Impact of knocking *HLTF* out on the repair profile produced at TKOv3 Cut site 2 in RPE1-*Cas9* TKOv3 cells. Assay performed as in figure 1C. Three independent *HLTF^-/-^* clones are shown. n = 3, except for clone 9 (n = 4), error bars show ±SD. **(C)** Effect of PCNA on the repair profile of wild type and *HLTF^-/-^* RPE1-*Cas9* TKOv3 cells. After transfection with NT or *PCNA*-targeting gRNAs, cells are arrested in G0/G1 by confluency for 12 days and then transfected with the TKOv3 Cut site 2 gRNA. Genomic DNA is purified three days later. n = 3 for WT + *NT* gRNA, n = 5 for WT + *PCNA* gRNA, n = 2 for *HLTF^-/-^* + *NT* gRNA and n = 5 for *HLTF^-/-^* + *PCNA* gRNA. Error bars show ±SD. **(D)** Repair pattern obtained at *AAVS1* cut site 4 in the RPE1-*Cas9 HLTF^-/-^* cell line complemented with the indicated versions of *HLTF*. Doxycycline is added to induce *HLTF* expression (figure S5C) 24 hours before transfection with *AAVS1* cut site 4 gRNA, and it is maintained until genomic DNA purification (3 days after transfection). Only most frequent indels are shown. n = 3 except for the RING mutant (n = 2). Error bars show ±SD. **(E)** ChIP q-PCR experiments showing HLTF binding to TKOv3 Cut site 2 in the indicated genotypes at 0 and 3 hours after transfection with TKOv3 Cut site 2 gRNA. *GREB1* locus was used as a control site. n = 3, error bars show ±SD. Statistical analysis was performed using Two-way ANOVA. **(F)** (left) ChIP q-PCR experiments showing Cas9 binding to TKOv3 Cut site 2 in the indicated genotypes at different time points after transfection with the TKOv3 Cut site 2 gRNA. *GREB1* locus was used as a control site. (right) Cas9 cleavage efficiency at TKOv3 Cut site 2 estimated by qPCR (as in figure 4B) in the same samples used for Cas9 ChIP experiments in wild type and *HLTF^-/-^*backgrounds. n = 3, error bars show ±SD. Statistical analysis was performed using Two-way ANOVA. **(G)** Experimental layout diagram for the in vitro removal of post-cleavage Cas9 RNP by HLTF (left, see figure S5F and Methods). Immobilized dsDNA, containing the TKOv3 Cut 2 gRNA target sequence, is digested by the addition of TKOv3 Cut 2 gRNA-Cas9 RNP. Retained Cas9 RNP is then released by incorporation of increasing bacteria-purified HLTF concentrations to the mix. (Right) Representative electrophoresis gel image of flow-through material after incubation with HLTF. Normalized quantification of the released Cas9 bands is shown below the electrophoresis image.

HLTF-mediated fork reversal involves protein removal and a rearrangement of nucleic acid strands (*20*) A comparable situation occurs in post-cleavage Cas9 RNP complexes, with Cas9 protein and the hybridized gRNA remaining bound to the DSB. Thus, HLTF could help remodel this structure by removing Cas9 and reannealing DNA ends for subsequent fill-in repair. Indeed, HLTF was recruited to Cut 2 target site 3 hours after transfection with its corresponding gRNA, suggesting a direct function in the DSB repair process (figure 5E). Furthermore, Cas9 protein accumulation at the target site following DSB induction was significantly increased in the absence of HLTF (figure 5F), without significant changes in cutting kinetics, indicative of a higher residence time of the Cas9 RNP when HLTF was not present. Finally, we tested whether HLTF had indeed the capability to remove the Cas9 RNP from the post-cleavage complex *in vitro* (figure 5G). For this, we assembled a DNA substrate onto streptavidin magnetic beads using biotinylated DNA. Next, we incubated pre-assembled Cas9-gRNA with the beads to load Cas9 onto the DNA substrate. After washing the excess of Cas9, beads were incubated at increasing concentrations of human HLTF and released Cas9 was analyzed on an SDS-PAGE gel by Coomasie staining or Cas9 immunodetection. Results, with two independent hHLTF purifications from *E. coli* (figure 5G) and insect cells (figure S5F), suggest that HLFT has the biochemical capacity to detach Cas9 from post-cleavage complexes. Taking these data together, we have identified HLTF as a novel human factor facilitating the removal of Cas9 RNPs from the post-cleavage complex, affecting thus subsequent processing and the repair outcome.

### The Fanconi pathway, and the BTRR and SAGA complexes are involved in MMEJ

*POLQ* was a selected hit in the screen, showing a repair pattern of decreased microhomology-mediated and bidirectional deletions (figure 1F), therefore compatible with a specific deficiency in MMEJ. This pattern was confirmed in two independent p53-proficient clones (figure 6A). However, the cluster corresponding to gene knockouts with a similar event distribution as *POLQ* deletion, and therefore also likely to be involved in MMEJ, was not as well-defined as those of NHEJ factors or nucleases (figure 2D). This is likely because, in the RPE1 cell line, NHEJ events are more prevalent, at least for the cut-sites tested, so the overall difference with the wild-type profile is minimized when only MMEJ is affected (e.g., distance: *POLQ* = 7.53; *POLL* = 53.79). We therefore selected 25 gene knockouts that resulted in an indel/event distribution similar to that of *POLQ* (PCC < 0.45), regardless of the overall differences with control patterns. STRING analysis of these genes showed a highly significant enrichment for functional interactions (p-value < 1.0e-16) (figure 6B), with the BLM-TOP3-RRMI1-RRMI2 (BTRR) complex (FDR = 0.0057), the SAGA chromatin-modifying complex (FDR = 5.86e-07) and the Fanconi anemia pathway (FDR = 3.29e-08). The repair pattern of representative mutants of these three pathways/complexes (*BLM*, *TADA1* and *FANCF*), which were, as a matter of fact, also among the 168 selected genes in the screen, was similar to that of *POLQ* deficiency (compare figure 6C and 1F, figure S6A). These data suggest previously uncharacterized functions of these pathways and complexes in MMEJ, while also validating the use of REPAIRome data mining to reveal novel repair functions and pathways simply based on the similarity between indel distributions.

**Fig. 6.**
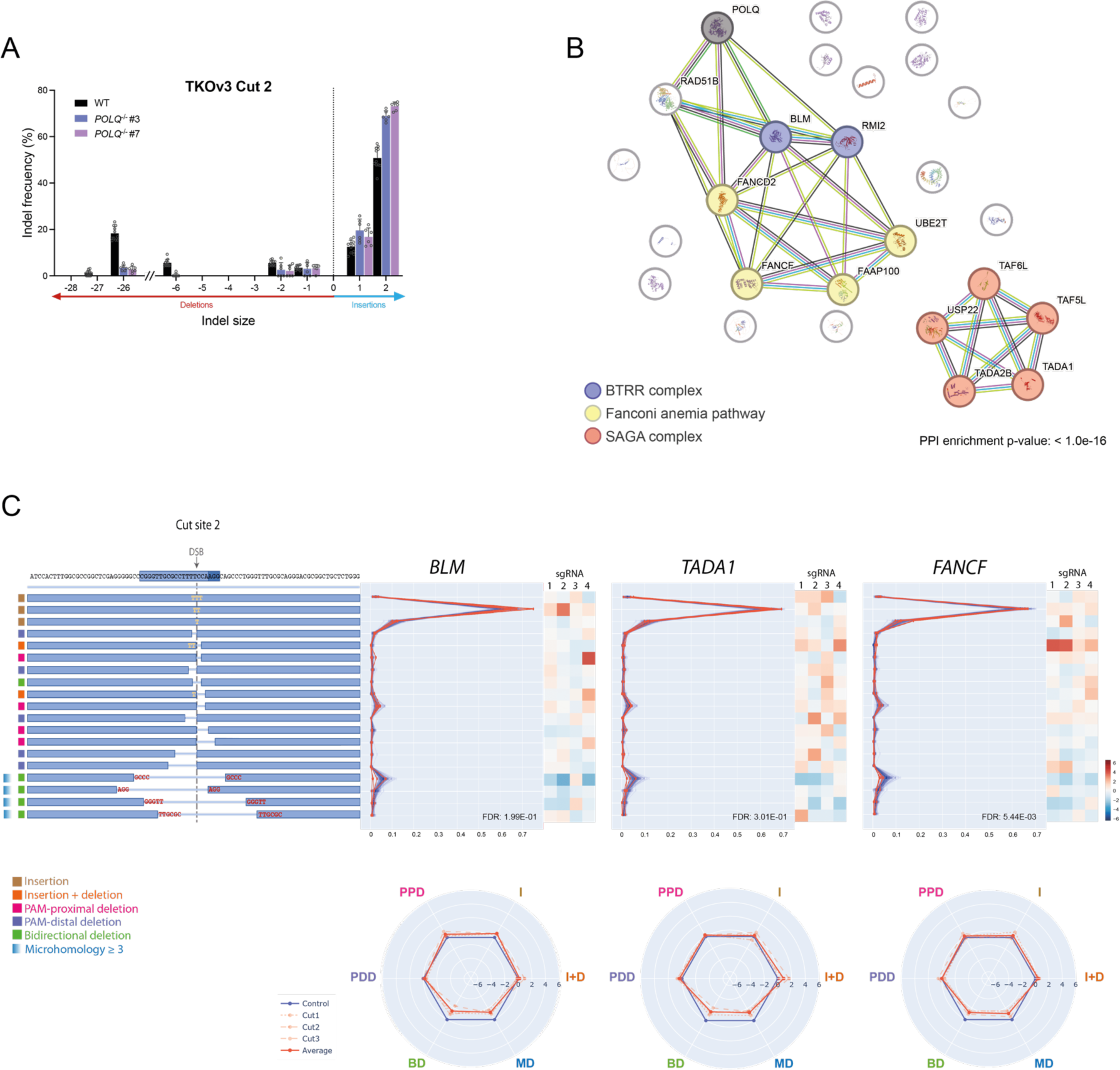
The BTRR complex, Fanconi pathway and SAGA complex are involved in MMEJ. **(A)** Impact of knocking *POLQ* out on the repair profile observed at TKOv3 Cut site 2 in RPE1-*Cas9* TKOv3 cells. Assay performed as in figure 1C. Two independent *POLQ^-/-^* clones are shown. n = 11 for wild type and 6 for *POLQ^-/-^*clones. Error bars show ±SD. **(B)** STRING analysis of interactions between selected proteins (i. e., POLQ PCC > 0.45). Only functionally connected genes are annotated. **(C)** Summary of screen results for cut site 2 for *BLM*, *TADA1* and *FANCF*. Displayed as in figure 1D.

### ID11 cancer mutational signature is caused by insertional NHEJ and favored upon VHL loss or hypoxia

EGLN1 and VHL are two key factors controlling the hypoxia response by, respectively, first hydroxylating and then ubiquitinating hypoxia-inducible factor (HIF)-α transcription factors, promoting their degradation in normoxic conditions. Upon hypoxia, HIF-α proteins are stabilized and trigger a hypoxia-adaptive transcriptional response. Both *EGLN1* and *VHL* were identified as factors significantly affecting the indel pattern, and clustered together with genes whose knockout pushed DSB repair towards insertions (figure 2C, 7A and S7A). We therefore selected *VHL* for further characterization. The DSB repair phenotype was confirmed in *VHL* pooled knock-out populations (see table S3 for knock-out efficiencies) with a mild, but significant, increase in insertions (figure 7B), most evident at the predominant +2 event of TKOv3 Cut 2. We were unable to recover individual *VHL^-/-^* clones in p53-proficient background. The phenotype conferred by *VHL* pooled knock-out was lost upon concomitant deletion of *HIF1A*, but not of *HIF2A* (figure 7D). In addition, the insertion-promoting behavior of VHL-deficient populations was observed, to a variable extent, in four endogenous *AAVS1* cut sites (figure 7C). Altogether, these results suggest that loss of *VHL* favors insertions through a mechanism involving the HIF1α-dependent hypoxia response.

**Fig. 7.**
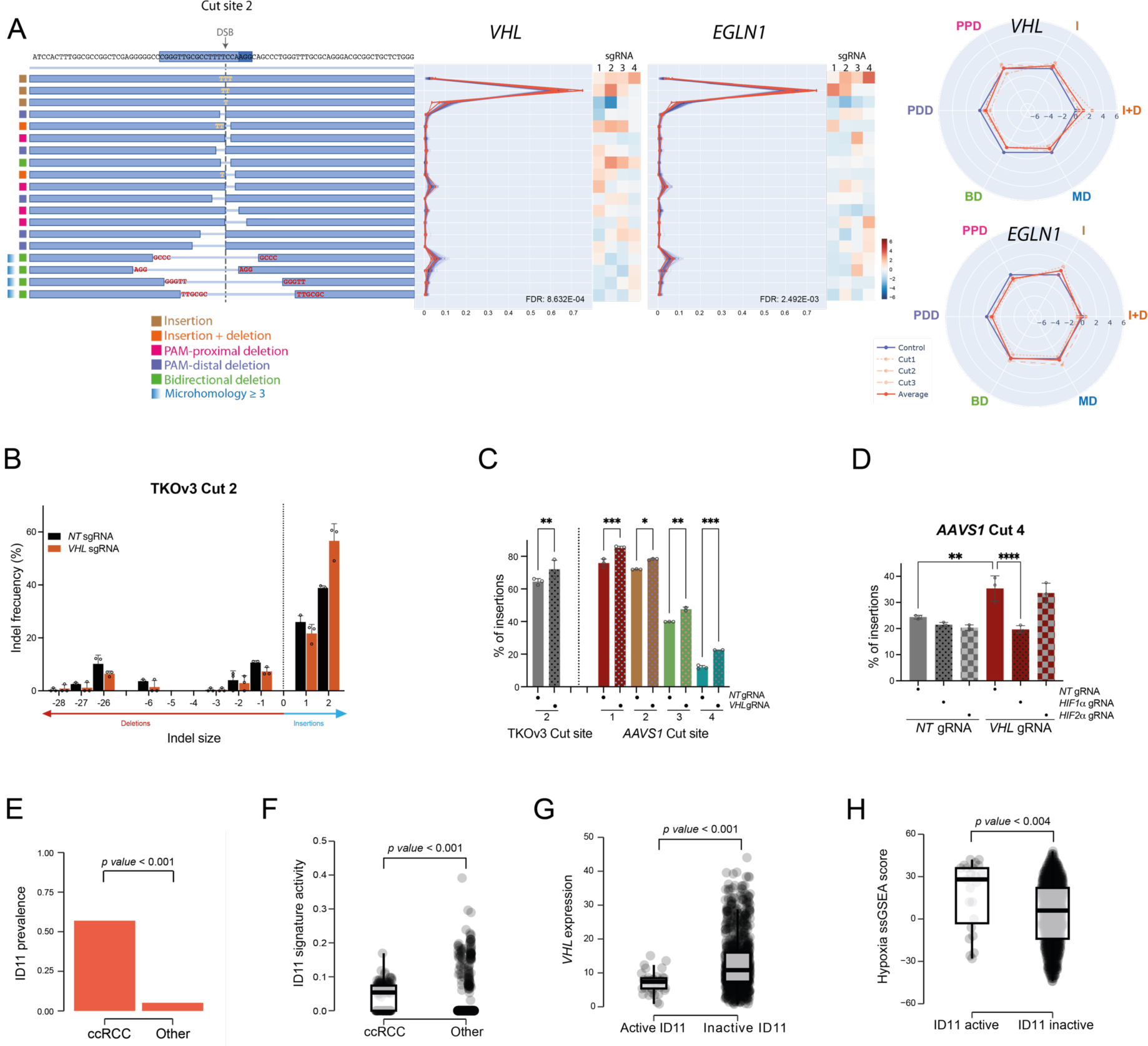
ID11 cancer mutational signature is caused by insertional NHEJ and favored upon VHL loss or hypoxia. **(A)** Summary of screening results for *VHL* and *EGLN1*, as in figure 1D. Only TKOv3 Cut 2 data is shown. **(B)** Effect of indicated *de-novo* designed gRNAs transfection on the repair profile produced at TKOv3 Cut site 2 in RPE1-*Cas9* TKOv3-NT cells. Assay carried out as in figure S3A (see also Methods). n = 3, error bars show ±SD. **(C)** Effect of the indicated gRNAs transfection on the percentage of insertions relative to all indels obtained at the TKOv3 Cut site 2 and at different endogenous *AAVS1* Cut sites. Assay carried out as in figure S3A. n = 3, error bars show ±SD. Statistical analysis was performed using One-way ANOVA. **(D)** Effect of the indicated gRNAs co-transfection on the percentage of insertions relative to all indels obtained at the *AAVS1* Cut site 4. Assay carried out as in figure S3A. n = 3, error bars show ±SD. Statistical analysis was performed using One-way ANOVA. **(E)** Comparison between the relative frequency (i.e., prevalence) of tumour samples with detectable ID11 signature activity in 144 ccRCC samples and 2,464 tumour samples from other 36 tumour types represented in PCAWG. P-value is calculated using Fisher’s exact test. **(F)** Comparison between ID11 signature activity in 144 ccRCC samples and 2,464 tumour samples from other 36 tumour types in PCAWG. P-value is calculated using unpaired, two-sided Wilcoxon test. **(G-H)** Comparison of *VHL* expression (FPKM counts) (G) and hypoxia scores (H), as previously estimated (*83*), using a reported signature (*84*), with activity of ID11 in 1,067 non-ccRCC tumour samples with available RNA-Seq expression in PCAWG. *VHL* expression and hypoxia scores in active ID11 (n = 37, ID11 signature activity > 0) and inactive ID11 (n = 1030, ID11 signature activity = 0) are compared using an unpaired, two-sided Wilcoxon test.

*VHL* is inactivated in virtually all cases of clear-cell renal cell carcinoma (ccRCC) (*57*), a type of cancer that is associated with a high load of indel mutations, for which the causative mechanism remains unknown (*58*). To address a possible connection between VHL loss and the characteristic indel mutational load of ccRCC, we checked the prevalence of the 17 indel mutational signatures (ID1-17) described in the COSMIC database (https://cancer.sanger.ac.uk/signatures/id) across 37 cancer types (*59*). ID11 appeared as the only indel signature that was particularly enriched in ccRCC when compared to other cancer types. Indeed, ccRCC was the only cancer with more than 50% prevalence of ID11 (figure S7B), and, when compared to an aggregate of all the remaining tumors, ccRCC showed a significant increase in both ID11 prevalence and activity (figure 7E and F). ID11 has a yet-unknown molecular etiology and is characterized by unbiased one-nucleotide insertions not driven by the pre-existence of an homopolymer (figure S7C), which is compatible with the pattern of insertional NHEJ events that are favored upon *VHL* loss. Further supporting the association between *VHL* and ID11, non-ccRCC tumors with active ID11 were significantly associated with lower expression of *VHL* and with a hypoxic transcriptional profile (figure 7G and H). We therefore propose that ID11 can result from insertional NHEJ events favored in hypoxia and the pseudo-hypoxia conditions that accompany *VHL* loss.

## Discussion

The REPAIRome resource presented here, with its interactive browsable webtool (will be accessible upon publication of the work), enables the analysis of the contribution of the majority of human genes to Cas9-induced DSB repair outcomes in a model human untransformed cell line. By searching for a given gene of interest, it readily returns key information to understand the effect of this gene on the repair profile such as the distance to controls, the distribution of main categories and sub-categories of events, as well as the complete indel distribution and fold-change for each sgRNA in the two replicates and three cut sites. This allows for an intuitive visualization of repair patterns, as well as a more complete evaluation of the significance of the results (see figure 1D-F for examples). The webtool also includes a table with the list of genes whose indel distribution correlates with that of the gene of interest (PCC ≥ 0.45) and their corresponding distance to controls, as well as a STRING analysis highlighting functional relationships among them. We believe this is a powerful resource for the scientific community, and especially for those interested in DSB repair and the biotechnological and medical use of CRISPR-Cas systems.

In addition to individual gene analysis, the data generated can be mined to identify pathways based on the similarity between repair patterns, highlighting its potential to drive discoveries in the DSB repair field. This database and webtool will continue to be updated, as we deepen our analysis with additional libraries, cut sites, cell lines and conditions. The results can be further integrated with the rich chemosensitivity data in Olivieri et al. (*14*), as the same cell line and lentiviral knock-out CRISPR library have been used, providing a comprehensive view of the cellular response to DSBs. Moreover, the improvements implemented to enhance the throughput capacity of massive DSB-repair profiling provide a methodological framework that can be exploited in future studies, as with Repair-seq (*60*, *61*), to explore novel mediators of base-or prime-editing in a genome-wide manner, or in more sophisticated setups such as combinatorial knock-out libraries to explore genetic relationships or saturation mutagenesis of particular genes of interest.

Our study provides several examples that serve as validation of the power of the resource generated. First, we provide evidence that challenges the current view of a collaborative/redundant function of XLF and PAXX paralogs in promoting end synapsis for ligation during NHEJ. Instead, we propose that PAXX affects the equilibrium of XLF-mediated synaptic structures so that nucleolytic processing is favored (figure 3C). This explains the screening results and genetic relationships observed in the context of known structural functions of these factors. While POLL loss only affects insertions, XLF deficiency also decreases short deletions in favor of longer events, further supporting this structural rather than enzymatic function. Our model is also consistent with the synergistic effects for *XLF* and *PAXX* mutations previously observed in animal model viability, DNA damage sensitivity and DSB repair (*32–37*, *62*), but with the two paralogs operating in alternative and competing end-joining sub-pathways. Thus, the molecular information provided by REPAIRome has uncovered novel mechanistic insight into the function of XLF and PAXX.

Second, we find that different components of the cellular endomembrane system can affect CRISPR-Cas9-mediated cutting efficiency, likely through an effect on the uptake, distribution and/or stability of the gRNA. In addition to relevant implications for gene editing strategies and the choice of delivery systems and routes, these results further uncover that cutting efficiency impacts on overall editing not only quantitatively, but also qualitatively, affecting the indel profile. At least in some cases, this is caused by re-cutting of already-edited targets, resulting in outcomes that are a composite of more than one mutagenic repair event, and which are consequently disfavored in conditions of low cleavage. This is particularly evident for +2 insertions, which arise as two consecutive +1 events. Current understanding is that +2 events result from variability in staggered cutting by Cas9 (*30*, *51*, *52*), but recent massive analysis of Cas9 cleavage has evidenced a wide majority of blunt and 1-nucleotide staggering, with longer overhangs only representing very rare events (*63*). We thus propose that multi-nucleotide insertions are generated by sequential templated single-nucleotide insertions after target re-cleavage, in agreement with previous observations in budding yeast (*29*). Understanding the tolerance of Cas9 RNP for nucleotide mispairing at the cut site is therefore important to predict indel outcomes and provides new grounds for improving current prediction software; e.g., inDelphi predicts predominant +1 insertions for Cut 2, instead of the +2 observed experimentally.

Third, we identify the tumor suppressor HLTF as a novel factor involved in the repair of Cas9-induced DSBs. Although the role of this protein in DNA damage tolerance and response to replicative stress is well established, a direct connection with DSBs has not been reported. There are, however, indications that the yeast homolog Rad5 is also involved in DSB repair (*64*), suggesting evolutionary conservation. We here show that, at least *in vitro,* HLTF can facilitate unloading of Cas9 RNP post-cleavage complexes, thus releasing free DNA ends. This activity is consistent with the increased residency of Cas9 at the cut site in *HLTF^-/-^*cells. HLTF may thus facilitate insertions by favoring the reannealing of gRNA-displaced strands for fill-in reactions. In a physiologic context the activity of HLTF may be relevant for the removal of DNA-RNA hybrids at DSBs, formed by either breakage of R-loop structures (*65*) or the reported *de novo* transcription at DSB sites (*66*). The exact mechanistic details of HLTF function in DSB repair, and why this seems to depend on the sequence context at the break site is an exciting avenue for future investigation.

Fourth, we identify the BTRR complex, Fanconi Anemia pathway and SAGA as relevant cellular factors affecting MMEJ. Importantly, we do so by analyzing gene knockouts with an indel distribution pattern similar to that of *POLQ*, demonstrating the potential of REPAIRome data mining for discovering relevant connections in DSB repair pathways. We do not find, however, classic MMEJ factors such as PARP1, XRCC1-LIG3 or XPF-ERCC1 (*67*) nor proteins that have been implicated more recently such as APEX2 or the 9-1-1 complex and RHINO (*68*, *69*). It is possible that the cell line and conditions used in the REPAIRome screen limit the detection of MMEJ events, which only represent between 2 and 7% (depending on the cut site) of all indels in control conditions. The MMEJ-deficient phenotype of BTRR and Fanconi factors may be caused by a defect in resection (*67*, *70–73*) or, alternatively, in additional steps of MMEJ such as microhomology pairing or POLQ-mediated DNA synthesis. The involvement of the SAGA complex in resection is less clear, as it has only been observed in yeast and in conditions in which the redundant function of the Nua4a acetyltransferase complex is lost (*74*). It is tempting to think that the function of SAGA in MMEJ is connected to the reported local transition in H2B-K120 histone marks from ubiquitination to acetylation upon the induction of DSBs (*75*), and whose function and impact on DSB repair remains enigmatic. The SAGA complex may be a druggable target to impair MMEJ, and therefore likely to sensitize homologous recombination-deficient tumors, representing an alternative to, or a possible combination with, POLQ inhibition in cancer treatment (*76*).

Finally, we uncover that *VHL* (and *EGLN1*) deletion favors insertions in a HIF1α dependent manner, indicating that hypoxia conditions rewire DSB repair pathways and affect repair outcomes. Connections of VHL and the hypoxia response with DSB repair have been previously established, mainly based on changes in the expression of DNA damage response genes, but the results remain somewhat controversial regarding specific effects on NHEJ and MMEJ (*77*). Most importantly, we have linked VHL deficiency, and hypoxic conditions in general, to the specific cancer mutational signature ID11. The molecular etiology of this signature remained unknown, but is perfectly compatible with NHEJ-mediated insertions, as the lack of repeated nucleotides eliminates the possibility of polymerase slippage as in ID1 and ID2 mutational signatures. We therefore propose that the molecular etiology of ID11 is insertional NHEJ, and that this is favored in hypoxic conditions or upon loss of the tumor suppressor VHL. This explains the high indel burden reported for ccRCC tumors (*58*), which are invariably defined by VHL deficiency (*57*). In addition to the intrinsic value of this discovery, it represents a proof of concept on how the REPAIRome resource and methodology can be applied to understand genome instability in physiologically relevant processes such as cancer. In this sense, deeper analysis, for example expanding the number of cut sites and repair events, could constitute the basis for a more comprehensive and unbiassed analysis of indel mutational signatures in the future.

Altogether, our results highlight the groundbreaking nature and potential of the REPAIRome resource as a tool to fuel discoveries on DSB repair mechanisms, CRISPR-Cas9-mediated gene editing, mutational signatures relevant in physiopathological processes and, ultimately, potential therapeutic interventions.

## Materials and methods

### Cell culture

All cell lines were cultured at 37°C 5% CO_2_ in standard tissue culture incubators. RPE1-hTERT Cas9 (RPE1, female human [*Homo sapiens*] retinal pigmented epithelium) and RPE1-hTERT Cas9 *TP53^-/-^* cells were a kind gift from Daniel Durocher’s laboratory. RPE1 cells were cultured in Dulbecco’s modified Eagle medium (DMEM:F12-Ham) (Sigma-Aldrich, #D8437). HEK293T cells (human [*Homo sapiens*] embryonic kidney) were originally obtained from the ATCC cell repository (CRL-3216) and cultured in DMEM (Gibco, #21090-022). All media were supplemented with 10% (vol/vol) fetal bovine serum (Gibco, #A5256701), 100 U ml^−1^ penicillin, 100 µg ml^−1^ streptomycin (Sigma-Aldrich, #P0781) and 2 mM L-glutamine (Gibco, #25030-081). Cell lines were routinely tested for mycoplasma.

### Cell line generation

Knockout cell lines were generated by transfection of the gRNA (12 nM, final concentration), targeting the gene of interest, using RNAiMAX (Invitrogen, #13778500) according to manufacturer’s instructions. For gRNA design we used both Synthego (https://design.synthego.com/<u>#/</u>) and CRISPOR (http://crispor.tefor.net/) (*78*) webtools and checked the candidate gRNAs KO generation predicted efficiency on InDelphi (https://indelphi.giffordlab.mit.edu) (*8*). Prior gRNA transfection, crRNA (containing the sequence targeting the gene of interest) was annealed with a tracrRNA to form a functional duplex. Both crRNA and tracrRNA were purchased from Integrated DNA Technologies. 48 hours after transfection, single cells were sorted into a 96 wells plate by using a BD Influx cell sorter (BD Biosciences). Clones were grown until reaching enough cell numbers and genetic knockouts were identified by PCR amplification, Sanger sequencing and TIDE analysis (http://shinyapps.datacurators.nl/tide) (*79*). Successful knockout generation was finally verified by immunoblot or RT-qPCR analysis. Sequences of gRNAs and primers for PCR amplification are listed in tables S7 and S8, respectively.

RPE1 Cas9 TKOv3-NT and RPE1 Cas9 *TP53^-/-^* TKOv3-NT cell lines were generated by lentiviral transduction using the TKOv3 library backbone plasmid (pLCKO2, Addgene #125518) with a non-targeting gRNA sequence cloned into the BsmB1 restriction sites. Lentivirus were produced by transfecting HEK293T cells with the pLCKO2-NT transfer vector together with the standard second-generation packaging vectors (pCMVdR8.74 and pMD2G) using Lipofectamine 3000 Transfection Reagent (Invitrogen, #L3000001). Viral supernatant was harvested 72 hours post-transfection and filtered through a 0.45 μm PVDF syringe filter (Millipore). Lentiviral infection was carried out at a low MOI (∼0.1 viral particle per cell) to achieve only one integration per genome. Transduced cells were selected by addition of puromycin (20 µg ml^−1^) (Merk, #P8833) to the culture medium.

Complementation of *HLTF^-/-^* cells lines with the different *HLTF* mutant constructs was performed by transducing the mutant *HLTF-*expressing lentiviral plasmids into the RPE Cas9 *HLTF^-/-^*cell line (clone #10). Lentiviral production and infection of the *HLTF^-/-^*knockout were carried out as described above. Transduced cells were selected by puromycin addition (20 µg ml^−1^) to culture medium. Inducible HLTF lentiviral vectors were a kind gift from Carlene Cimprich’s laboratory (*22*). To induce *HLTF* expression 20 nM doxycycline was added to the medium 24 hours before each experiment. We noticed that HLTF-DEAA levels after doxycycline-induction were notably lower than those observed for the rest of constructs. Therefore, doxycycline dose used to induce expression of this mutant *HLTF* was increased up to 1 μM (see figure S5C).

### Library representation and experimental depth determination with barcoded library – Cell culture and DSB induction

RPE1-hTERT Cas9 *TP53^−/−^* cells were transduced with the lentiviral CloneTrackerXP barcode library (50 M barcodes, Cellecta, #BCXP50M3RP-P) at a very low MOI to ensure only one integration per cell. After selection with puromycin, around a thousand cells were seeded on a separate dish and kept growing in the presence of puromycin. Then, transduced cells were divided into two technical replicates. For each replicate, a million cells were transfected independently with either CloneTracker_gRNA1 and CloneTracker_gRNA2 (table S7) as described above (see Cell line generation). Three days later, all cells were collected, pelleted and stored at −80 °C until Illumina library preparation.

### Library representation and experimental depth determination with barcoded library – Sequencing library preparation

To sequence the DSB repair products associated to each CloneTracker bracode, genomic DNA was extracted using DNeasy Blood and Tissue Kit (Qiagen, #69504). For library preparation, the lentivector region including both the barcode and Cas9 target site was amplified in two rounds of PCR, using primers CellTracker_P1Fwd and CellTracker_P1Rev (table S6). For the first round, four PCR reactions containing 2 μg of genomic DNA (gDNA) in 50 μl total volume were performed. These reactions contained amplification primers at a final concentration of 0.3 μM each, and 1X Q5 Mastermix Next Ultra II (New England Biolabs, #M0544L), and were run with the following program: 1 step of 3 min at 95 °C; 18 cycles of 30 s at 95 °C, followed by 30 s at 64°C and 30 s at 72 °C; and a final step of 2 min at 72 °C. For including Illumina adapters and indexes, 5 μl of pooled PCR products of each sample were used as a template for a second round of PCR to add Illumina P5 and P7 adaptors and i7 indexes. These PCR reactions were assembled in 100 μl reactions of 1X Q5 Mastermix Next Ultra II with primers at 1 μM final concentration. This PCR mix was split into x2 50 μl PCR reactions and run in a thermocycler with the following program: 1 step of 3 min at 95 °C; 11 cycles of 30 s at 95 °C, followed by 30 s at 67 °C and 30 s at 72° C; and a final step of 2 min at 72 °C. PCR products were purified with QIAquick PCR purification kit (Qiagen, #28106), followed by gel-purification using QIAquick gel extraction kit (Qiagen, #28704) and a last purification step using QIAquick PCR purification kit once more time. The sequence of all primers used for this section are listed in table S6.

### REPAIRome screen – TKOv3 CRISPR KO library amplification and lentiviral packing

The Toronto KO CRISPR library v3 (TKOv3) (*13*) was acquired from Addgene (#125517). For library amplification, four 50 μl aliquots of high efficiency electrocompetent *E. coli* cells (MegaX DH10B T1^R^, Thermofisher, #C640003) were electroporated (2.0 kV, 200 Ω, 25 μF) with 50 ng of the library plasmid in 0.1 cm cuvettes, following manufacturer recommendations. After recovery, serial dilutions of electroporated bacteria were seeded on LB plates with antibiotic selection to estimate transformation efficiency. The rest of bacteria were grown in 500 ml of LB with antibiotic overnight. After plasmid maxiprep (PureLink Maxiprep kit, Invitrogen, #K210007), original and amplified TKOv3 libraries were Illumina-sequenced to verify that sgRNA representation was maintained.

To generate lentiviral particles, HEK293T cells were transfected with an equimolar mixture of the lentiviral library DNA plus the packaging capsids psPax2 and pMD2.G using GeneJuice (Sigma-Aldrich, #70967). The culture medium was replaced 16 hours later with fresh D-MEM medium supplemented with 10% FBS, DNase I (1 U ml^−1^, Zymo Research, #E1010), MgCl2 (5 mM) and 20 mM HEPES pH 7.4 to prevent carryover of plasmid DNA into the viral preparation. Cell supernatants were harvested 72 hours post transfection, filtered through a Steriflip 0.45 µm filter (Merck Millipore, #SE1M003M00) and concentrated by ultracentrifugation at 10 000 rpm, 4 °C for 1 hour in a SW-32 rotor 1 hour after addition of 5 µg ml^−1^ of LentiFuge viral concentration reagent (Cellecta, #LFVC1). Viral pellets were resuspended in cold PBS with 10% FBS. For titration, 5 x 10^4^ HEK293T cells were infected with serial dilutions of the viruses. After 16 hours the medium was replaced with fresh medium and fluorescence was measured by FACS 48 hours later. Transducing units per milliliter were calculated from those dilutions that yielded between 1% and 20% of GFP positive cells.

### REPAIRome screen – Cell culture and DSB induction

RPE1-hTERT Cas9 *TP53^−/−^* cells were transduced with the lentiviral TKOv3 library as previously described (80). Briefly, RPE1-hTERT Cas9 *TP53^−/−^* cells were transduced with the lentiviral TKOv3 library at low MOI (∼0.3 −0.4). Two days later, 20 μg ml^−1^ puromycin was added to the culture medium to select for transductants. The next day cells were trypsinized and replated in the same plates while maintaining the puromycin selection to accelerate selection. The following day, which was 4 days after infection, was considered the initial time point (T0) and cells were pooled together and divided into two technical replicates (labeled as “A” and “B”). A pellet of 50 x 10^6^ cells was also stored at −80 °C for estimating library representation at the starting point of the screening. Replicates were then cultured independently for 12 days to allow for protein depletion while always maintaining at least 4.5 x 10^7^ cells per replicate after each pass to ensure library representation of >600-fold throughout the whole screening. At day 12 after infection, for each replicate 1 x 10^8^ cells (1200-fold library representation) were transfected independently with each of the three library-targeting gRNAs and 4.5 x 10^7^ (500-fold library representation) cells were transfected with a non-targeting gRNA (labelled as “T16 A-NT” and “T16 B-NT”). Transfections were carried out as described above (see Cell line generation). 4 days after transfection (T16), cells were pelleted and stored at −80 °C until Illumina sequencing library construction.

### REPAIRome screen – Sequencing library preparation

Pellets from T0 and from T16-NT (replicate A and B) were used to generate Illumina sequencing libraries to estimate library representation at starting (T0) and endpoint (T16). gDNA was extracted using QIAamp Blood Maxi Kit (Qiagen, #51194). Genome-integrated sgRNA sequences of 50 x 10^6^ cells per sample were amplified by multiple PCR reactions using primers TKOv3_P1_Fwd and TKOv3_P1_Rev. First round PCR reactions contained 3 μg of gDNA into 50 μl of total volume. These reactions contained amplification primers, at a final concentration of 1 μM each, and 1X Q5 Mastermix Next Ultra II, and were run on a thermocycler with the following program: 1 step of 3 min at 95 °C; 19 cycles of 30 s at 95 °C, followed by 30 s at 66 °C and 30 s at 72 °C; and a final step of 2 min at 72 °C. 5 μl of pooled PCR products of each sample were used as a template for a second round of PCR to add Illumina P5 and P7 adaptors and i7 indexes. These PCR reactions were assembled in 200 μl reactions of 1X Q5 Mastermix Next Ultra II with primers at 1 μM final concentration. This PCR mix was split into x4 50 μl PCR reactions and run in a thermocycler with the following program: 1 step of 3 min at 95 °C; 12 cycles of 30 s at 95 °C, followed by 30 s at 66 °C and 30 s at 72 °C; and a final step of 2 min at 72 °C. PCR products were purified as described for CloneTracker library preparation. The sequence of all primers used for this section are listed in table S6.

To sequence repair product generated at the end point (T16) of the REPAIRome screen, genomic DNA of all T16 samples transfected with each library-targeting gRNA was extracted using DNAzol (Invitrogen, #10503027) following manufacturer protocol, due to its higher yield compared to column-based extracting method. Then, the library lentivector region containing both the sgRNA and the DSB repair outcomes of 1 x 10^8^ cells per sample was amplified by multiple PCR reactions using 3.5 μg of gDNA in 50 μl, total volume, of 1X Q5 Next Ultra II Mastermix with primers TKOv3_P1_Fwd and RPAIRome_Rev_V2 at a final concentration of 1 μM. These PCR reactions were run with the same conditions as used for T0 and T16-NT samples. PCR products were pooled, and 200 μl of the pool were purified by size selection using Sera-Mag Select Beads (Cytiva, #11548692) at 0.55X, keeping the supernatant, followed by a second round at 0.8X, eluting the DNA in 100 μl of nuclease-free water. To incorporate Illumina adapters and indexes, 10 μl of purified PCR products of each sample were used as a template for a second round of PCR using staggered primers to add Illumina P5 and P7 adaptors as well as i5 and i7 indexes. These PCR reactions were assembled in 200 μl reactions of 1X Q5 Mastermix Next Ultra II with primers at 1 μM final concentration. This PCR mix was split into x4 50 μl PCR reactions and run in a thermocycler with the following program: 1 step of 1 min at 98 °C; 10 cycles of 10 s at 98 °C, followed by 30 s at 67 °C and 30 s at 72 °C; and a final step of 2 min at 72 °C. PCR products were purified as described for CloneTracker library preparation. The sequence of all primers used for this section are listed in table S6.

### Illumina Sequencing

Sequencing libraries from CloneTracker experiment, REPAIRome screen library representation, and T16 replicate B-Cut site 2 were sequenced on an Illumina NextSeq550 system. CloneTracker and REPAIRome screen library representation libraries were sequenced with a total of 2 reads per cluster (single read plus one index read) with the following read lengths: I1 = 6 nucleotides (sample index); R1 = 85 nts (Library barcode plus DSB repair outcome). T16 replicate B-Cut site 2 was sequenced with a total of 4 reads per cluster, including paired end reads with the following read length: I1 = 6 nts (sample index); I2 = 8 nts; R1 = 22 nts (dark cycles) + 21 nts (TKOv3 sgRNA); R2 = 248 nts (DSB repair outcome).

The remaining sequencing libraries from REPAIRome screen samples (T16), including Ca9 cut sites 1, 2 and 3 for replicate A; and cut sites 1 and 3 for replicate B, were sequenced on an Illumina Novaseq 6000 System with a total of 4 reads per cluster, including paired end reads plus two index reads with the following length: I1 = 8 nts (sample index i5); I2 = 8 nts (sample index i7); R1 = 22 nts (dark cycles) + 21 nts (TKOv3 sgRNA); R2 = 241 nts (DSB repair outcome).

### Focused DSB repair profile experiments

To study DSB repair outcome distribution in the different genetic backgrounds, knockout cell lines were transfected with a gRNA targeting the locus of interest (either the TKOv3-nt backbone or endogenous *AAVS1* locus) as described above (see Cell line generation), and 3 days later, cells were collected and gDNA was extracted with Qiagen DNeasy Blood and Tissue kit. Then, the genomic region including each target site was amplified by PCR using GoTaq Hot Start polymerase (Promega, # M5001). PCR reactions contained 400 ng of gDNA and 0.75 μM of the corresponding primers in a total volume of 30 μl. PCR products were then purified using QIAquick PCR purification kit and sequenced with ABI 3730xl sequencer (ThermoFisher, #A41046) using the corresponding sequencing primer. Finally, sequencing data was analyzed using the Sanger trace deconvoluting software Synthego ICE CRISPR Analysis Tool v3.0 (https://ice.synthego.com/).

In the case of polyclonal knockout pools, cells were transfected with the corresponding gene-targeting gRNA as described above (See Cell line generation) and cultured for 12-14 days to allow for protein depletion. These polyclonal knockout populations were then transfected with a second gRNA targeting the indicated site, either on the TKOv3 backbone or on the *AAVS1* locus, to generate the DSB used to analyze the repair outcomes. Polyclonal knockout efficiency was checked at the experimental endpoint by PCR amplification, Sanger sequencing and trace deconvolution analysis as described above.

Sequences of DSB induction gRNAs used for these experiments and sequences of gene-targeting gRNAs are listed in table S7, while sequences of primers used in this section are listed in table S9.

### Immunoblots

Whole-cell lysates were prepared by boiling the cells with Laemmli/SDS buffer 2X (120 mM Tris-HCl pH 6.8, 4% SDS, 20% glycerol). Protein concentration was measured by Nanodrop and equalized prior addition of bromophenol blue (0.01% w/v) and β-Mercaptoethanol (10% v/v). Approximately 30-40 μg of protein were loaded onto either 10% in-house acrylamide gels or 4-20% Mini-PROTEAN Tris-Glycine Precast Protein Gels (BioRad, #4561094). Separated proteins were transferred to Odyssey nitrocellulose membranes (LI-COR Biosciences, #926-31092); immunoblotted with the indicated antibodies and subjected to analysis using the Odyssey CLx system and ImageStudio Odyssey CLx software (LI-COR Biosciences). Primary antibodies: XLF (1:2000, TBST 5% Milk, Bethyl A300-730, Rabbit), PAXX (1:750, TBST 5% Milk, Abcam AB 126353, Rabbit), POLL (1:500, TBST 5% Milk, Bethyl A301-640A, Rabbit), Cas9 (1:1000, TBST 1% BSA, Abcam ab271303, Rat), Tubulin (1:10000 TBST 1% BSA, Sigma T9026, mouse) HLTF (1:1000, TBST 1% BSA, Abcam AB183042. Rabbit), PCNA (1:1000, TBST 1% BSA, Santa Cruz sc-56).

### Multi-site gRNA experiment

RPE1 Cas9 TKOv3-NT and *GET3^-/-^* clones #2 and #7 were transfected with 1.5 nM of a multi-site gRNA previously reported to target multiple sites in the genome (*50*) (table S7) as described above (see Cell line generation). 12 hours after transfection, cells were fixed with 4% paraformaldehyde for 10 min and processed for immunofluorescence. Samples were permeabilized with PBS 0.5% Triton for 15 min, blocked in 5% BSA in PBS and stained with anti-53BP1 primary antibody (1:500, 5% BSA PBS, Novus NB100-304, Rabbit) for 1 hour at room temperature, and secondary antibody coupled to Alexa Fluor 488 (1:1000, 5% BSA PBS, Invitrogen A11008, goat). DAPI counterstaining and ProLong mounting (ThermoFisher, #P36930) were performed for image acquisition using a LEICA confocal microscope SP5. The number of 53BP1 foci was evaluated in DAPI-selected nuclei using CellProfiler 4.2.1 analysis software.

### In-vitro Cas9 digestion

The constructs used for this assay were generated by PCR amplification, using Q5 Mastermix Next Ultra II, of 10 ng of either WT or +1 gBlock (Integrated DNA Technologies) as template and gBlock primers Fwd and Rev at a final concentration of 500 nM in a thermocycler with the following settings: 1 step of 1 min at 98 °C; 30 cycles of 10 s at 98 °C, followed by 30 s at 58 °C and 30 s at 72 °C; and a final step of 2 min at 72 °C. PCR products were purified with QIAquick PCR purification kit.

For the in-vitro digestion, Alt-R® *S.p.* Cas9 Nuclease V3 (Integrated DNA Technologies, #1081058) was used following a standard protocol. Briefly, Cas9-gRNA ribonucleoparticles (RNPs) were assembled prior to digestion by mixing 30 pmol Cas9 and 30 pmol gRNA in PBS in a total volume of 8.57 μl and incubating it for 15 min at RT. 2 μl of this RNP mix was added to a digestion mix containing 100 ng of purified PCR product (around 0.35 pmol) in a total volume of 15 μl of Cas9 cleavage buffer for the 10:1 RNP:oligo ratio. For the 5:1 and 2.5:1, subsequent 2-fold dilutions of the original RNP mix were performed to maintain the reaction volumes described above. A non-RNP negative control was included, where 2 μl of PBS were added instead of RNP. The DNA constructs were digested by incubating the reactions at 37 °C for 1 hour. Subsequently, the Cas9 protein was eliminated by incubating at 56 °C for 30 min with Proteinase K (1.33 mg ml^-1^). Finally, the complete reaction volumes were run in a 2% agarose TBE electrophoresis gels and visualized using a Biorad UV analyzer. The sequences of the gBlocks and primers used in this section are listed in table S10.

### RT-qPCR

Total RNA was extracted using the RNeasy Mini Kit (Qiagen, #74104) and genomic DNA was degraded using RQ1 DNAse (Promega, #M6101) following manufacturer’s protocol. 1 µg of nanodrop-quantified total RNA was used for cDNA synthesis with the Maxima H minus First strand cDNA synthesizer kit (ThemoFisher, #K1652) using random hexamer primers and standard protocol conditions in a final volume of 20 μl. The RT reaction was performed in three steps: 10 min 25 °C, 30 min 50 °C and 5 min 85 °C. The RT reaction was diluted 4.5-fold and 4.5 μl were used for the qPCR reaction. qPCR was performed using Syber-green 2X mix (Applied Biosystems, #4368577) in 10 μl reactions with gene-specific primers at a final concentration of 0.5 μM. All reactions were performed in technical triplicates and data were normalized to *GAPDH* levels. The sequence of the primers used in this section are listed in table S8.

### Cas9 cutting efficiency determination

Cells were transfected with TKOv3 Cut 2 gRNA as described previously (see Cell line generation). 6 hours after transfection cells were collected and genomic DNA was extracted using Qiagen DNeasy Blood and Tissue kit. Then, 50 ng of nanodrop-quantified gDNA was used for qPCR as described above (see RT-qPCR). All reactions were performed in technical triplicates and cutting efficiencies were normalized to the levels of amplification of the genomic locus *GREB1.* The sequence of the primers used in this section are listed in table S9.

### ChIP-qPCR

Cells on a 10 cm culture dish (at 70 – 80% confluency) were crosslinked with 1% formaldehyde in culture medium for 10 min at 37 °C. 125 mM glycine was then added to quench the crosslinking reaction and dishes were washed twice with ice-cold PBS. Subsequently, cells were scrapped in ice-cold PBS with Complete Protease Inhibitor Cocktail (Roche, #P1860) and 1 mM PMSF. After centrifugation (300 xg, 5 min, 4 °C), cells were lysed as follows. First, cells were incubated for 10 min in 1 ml of Lysis Buffer A (5 mM Pipes pH 8, 85 mM KCl, 0.5% NP40, 1 mM PMSF, 1x Complete Protease Inhibitor) to isolate nuclei. Next, nuclei were centrifuged (1500 xg, 5 min) and resuspended in Lysis Buffer B (50 mM Tris HCl pH 8.1, 1% SDS, 10 mM EDTA, 1 mM PMSF, 1x Complete Protease Inhibitor Cocktail). Chromatin was then sheared by sonication by using a Bioruptor (Diagenode, UCD-200) at high intensity, with 25 cycles (30 seconds sonication followed by a 30-second pause) and clarified by centrifugation (17000 xg, 10 min, 4 °C).

To evaluate chromatin fragmentation, first, crosslinking was reverted by incubating 5% of chromatin volume in Lysis Buffer B with 0.25 mg ml^-1^ Proteinase K (AppliChem, #A3830) for 16 hours at 65 °C. DNA was purified by Phenol:cloroform extraction and ethanol precipitation and quantified with Nanodrop one. Fragment size distribution was considered acceptable if the smear observed in an agarose gel electrophoresis ranged from 300 to 600 bp.

For each immunoprecipitation, 30 μg of chromatin and 4 μg of the specific antibody were incubated in IP buffer (20 mM Tris-HCl pH 8, 150 mM NaCl, 0.1% SDS, 1% TritonX-100, 2 mM EDTA, 1x Complete protease inhibitor cocktail, 1 mM PMSF) overnight at 4 °C. Next, 25 μl of protein A and protein G Dynabeads (ThermoFisher, #10015D), previously blocked with BSA, were added and incubated with chromatin (4h, 4 °C). Beads were then sequentially washed with IP buffer, IP buffer with increased salt (500 mM NaCl), LiCl buffer (20 mM Tris-HCl pH 8, 0.25 M LiCl, 1% NP40, 1% NaDoc, 1 mM EDTA), and TE buffer. DNA was eluted from the beads in 100 μl of elution buffer (1% SDS, 100 mM NaHCO3) at 50 °C. Crosslinking was reverted by adding NaCl (200 mM) and Proteinase K (100 μg) and incubating overnight at 65 °C. Samples were treated with RNAseA (0.5 mg ml^-1^) and purified with QIAquick PCR purification kit. Finally, samples were subjected to qPCR as described above (see RT-qPCR), using the indicated primers (table S9).

### Recombinant protein expression and purification

#### Expression and purification of hHLTF from bacteria

hHLTF gene was cloned into a pRSF plasmid using IVA cloning (*81*). *E. coli* BL21(DE3) competent Cells (Thermo, #EC0114) were transformed with plasmid pRSF-10xHis-TEVsite-hHLTF and grown in LB medium at 37 °C. Protein expression was induced with 1 mM IPTG when the optical density reached 0.6. The temperature was then lowered to 20 °C, and the bacteria pellet was collected after 16 hours. The bacterial pellet was lysed using a Spex SamplePrep 6875 Freezer/Mill in lysis buffer (25 mM Tris pH 7.5, 300 mM NaCl, 5 mM MgCl_2_, 10 mM Imidazol, 10% Glycerol). After lysate centrifugation at 50,000 xg for 45 min, the supernatant was passed through an affinity column (HisTrap HP, Cytiva, #17524802), and hHLTF was eluted with 250 mM of Imidazole. Next, hHLTF fractions were loaded on a cation exchange column (HiTrap-Q, Cytiva, #17115301) and eluted in high salt buffer (25 mM Tris pH 7.5, 500 mM NaCl, 5 mM MgCl2, 10% glycerol). Purified hHLTF was concentrated using Amicon Ultracentrifugal Filter (Merck, #UFC903008) and injected in Superdex 200 Increase 10/300 column (Cytiva, #28990944) (25 mM Tris pH 7.5, 200 mM NaCl, 5 mM MgCl2, 2 mM DTT, 10% Glycerol). The final purified protein was stored at −80 °C for future use.

#### Expression and purification of Cas9

Cas9 (pMJ915-spCas9) was expressed in Rosetta pLysS cells (Merck, #70956-M) and purified using a HisTrap column (20 mM Tris pH 8, 250 mM NaCl, 10 mM Imidazole, 10% Glycerol) with elution at 500 mM imidazole. The buffer was exchanged by dialysis (20 mM Hepes pH7.5, 150 mM KCl, 1 mM DTT, 10% glycerol), and the tag was removed by TEV protease digestion. An inverse HisTrap purification was performed to collect the cleaved Cas9 in the flow-through, followed by purification with an anion exchange column, where Cas9 was eluted with a gradient of increasing KCl concentration. Cas9 fractions were concentrated by Amicon Ultracentrifugal Filter and injected into a Superdex 200 Increase 10/300 column (20 mM Hepes pH 7.5, 150 mM KCl, 1 mM DTT). Purified Cas9 was stored at − 80 °C.

### HLTF release of Cas9 from DNA

To test HLTF’s ability to remove Cas9 from post-cleavage DNA complexes, 10 µL of Cas9 at a concentration of 1.3 µM per reaction tube were incubated with the corresponding gRNA targeting TKOv3 site 2 to assemble Cas9 RNP. Next, 10 µL of RNP were added to 10 µL of Dynabeads MyOne Streptavidin T1 beads (Invitrogen, #65601) with TKOv3_WT_gblock DNA substate immobilized in buffer-1 (25 mM Tris pH 7.5, 150 mM NaCl, 5 mM MgCl2, 2 mM DTT). Once the complex was formed, beads were washed three times with 30 µL of buffer-1. Next, beads were split into different reaction tubes (30 µL each tube) and incubated with increasing concentrations of HLTF (see figure S5F) in a final volume of 60 µL. Supernatant (flow-through) and remaining beads (beads) were collected for further analysis. To analyze the release of Cas9 by HLTF, samples (Flow-through, wash, beads) were run on SDS-PAGE and stained with Coomassie. When MBP-HLTF protein was used, western blotting with Anti-Cas9 antibody was used to detect Cas9.

### Analysis of cryo-EM particle distribution

To evaluate the effect of PAXX and XLF on the distribution of particles across the open and closed conformations of long-range DNA-PK complexes, we analyzed the metadata reported from several published cryo-EM studies F(*36*, *37*, *46–48*). Studies with insufficient detailed descriptions of data processing procedures or those that used modified substrates to stabilize particular conformations were removed from the analysis. Particle distribution is reported as the percentage of the total long-range complex particles found in those datasets. A summary of this data can be found in table S4.

### Analysis of representation and depth requirements in barcoded library

Fastq files with 80-nt single-end sequenced reads were quality-checked using FastQC and divided into two 40-nt fastq files. The first one contained the integrated barcode, and the second one contained the sequence of the repair outcome. The second file was aligned against the original sequence using Bowtie2, and then an indel profile was extracted for each barcode. This profile represented the percentage of reads containing an insertion/deletion based on the size of the indel, disregarding the position where it occurs.

To perform error simulation, we selected an increasing number of random barcodes with varying numbers of reads associated with them. Subsequently, we calculated the indel profile distribution corresponding to that number of barcodes and reads. For each combination, 100 simulations were conducted and compared against the mean indel distribution of all barcodes with more than 500 reads.

### Analysis of sequencing data from REPAIRome screen

Paired reads were sequenced to perform the analysis. The first read contained the information of the guide; to extract it, a pseudo-genome was created with each guide as a chromosome. This genome used the guide sequences from the TKOv3 library, extending the 3’ and 5’ ends with the vector sequence where it was integrated. Subsequently, these reads were aligned using Bowtie2 against the pseudo-genome to extract the name and the integrated guide in each read.

On the other hand, the second read contained information about how the cut was repaired. To extract this repair outcome, a custom python script based on ScarMapper software (https://github.com/Gaorav-Gupta-Lab/ScarMapper) (*82*) was employed. We used the same parameters, requiring a 10-nucleotide match upstream and downstream of the DSB, allowing only one mismatch in the sequence.

For each cut and replicate, we counted the number of different outcomes resulting from each analysis. Only outcomes appearing in 3 guides from the same gene/control were considered for subsequent analysis, resulting in hundreds to thousands of repair outcomes per replicate. Additionally, to be able to compare gene profiles, we required a minimum number of reads for the events we wanted to analyze. Thus, only events where 70% of the guides had at least two reads were used (80% in the case of cut 3 due to its greater diversity). Finally, common events in both replicates of each cut were selected, resulting in 17, 19, and 33 events for cut1, cut2, and cut3, respectively. In all the cuts, the most common events were conserved and were present in similar percentages in the two replicates, due to the reproducibility of the results. Finally, this analysis could potentially overlook events specific to a particular gene non present in 70% of all possible guides. To test this, we used DESeq2 for each cut to perform differential expression analysis, comparing the number of reads in the guides of all the proteins individually against the control guides. No new events resulted from this analysis.

In each sample, guides with fewer than 500 reads were considered unreliable and excluded, as were genes with fewer than three guides due to the aforementioned condition. Two normalizations were performed independently for each replicate of each cut. First, we calculated the frequency of an event for each guide by dividing the number of reads of that event by the total number of reads belonging to the guide. This normalization also included the “non-edited” event, which measures the number of reads without any indels. Then, a min-max scaler was used to normalize all variables, and the mean of the control guides was subtracted from the values. After this, the values of the two replicates were appended. Finally, for each gene, we took the median value of the guides, referred to as “absolute event distribution” in the paper.

Conversely, the same normalization was performed, but instead of using the total number of reads to divide the values for each event, we used only the sum of the reads containing indels and, therefore, excluding the non-edited reads. This frequency measures the relative distribution of one indel compared to the others. Using this frequency, we checked for differences in event distribution between one protein and controls, using the “Compositional” R package, allowing comparison of the relative distribution of the events of guides for one gene to the ones extracted from the 146 control sgRNAs. This test gave us a p value, that was corrected later using FDR. Later on, using also this frequency, we performed the same normalization as before, using min-max scaling, subtracting the mean of the control guides, appending the replicates, and taking the median values of each gene resulting in the “relative indel distribution” data. For each gene we have estimated the variable “distance” as the total Euclidean distance between that gene and the mean of the controls (including all events from the three cuts) using the relative indel distribution data.

A gene was selected if it met the following criteria: for at least one cut, the FDR was ≤ 0.05 for both replicates and the correlation between the relative indel distributions for both replicates was greater than 0.3. Finally, the distance between the gene and control sgRNAs had to be greater than 5.

To calculate the values for the radar plot, events were first classified into categories based on event type. A deletion was considered as “Microhomology deletion” when the homology at the site of the deletion was greater than three. Then, for each replicate, we added the percentages from the events belonging to the same category (without using the non-edited values to calculate the percentages). Finally, the means of the controls were subtracted, and the two replicates were appended. The values for each cut and gene were calculated as the median values of the two replicates appended. In these plots, genes were not filtered based on the number of guides. The “Editing efficiency” and the “Guide abundance” variables were normalized independently, using the number of reads edited divided by the total number of reads for the first one and the total number of reads for the second one. Then, the same steps as before were taken to normalize the values (min-max scaler…).

## Supporting information

Auxiliary Supplementary table 1

Auxiliary Supplementary table 2

Auxiliary Supplementary table 4

Auxiliary Supplementary table 6

Auxiliary Supplementary table 10

## Acknowledgments

We thank the Lumir Krejci laboratory for the kind gift of recombinant HLTF proteins, the CNIO Cytogenetics and Genomics Units, and Oskar Fernández-Capetillo for critical reading of the manuscript and suggestions. All authors except J.A.B. are hosted by the Centro Nacional de Investigaciones Oncológicas (CNIO), which is supported by the Instituto de Salud Carlos III and recognized as a ‘Severo Ochoa’ Centre of Excellence (CEX2019-000891-S) by MCIN/AEI/10.13039/501100011033.

## Author contributions

Conceptualization: FCL

Methodology: ELdA, IS, DGL, AFS, ECP, GM, RFL

Investigation: ELdA, IS, DGL, AFS, ECP, JTB, JBG, JAB, RFL

Funding acquisition: JAB, GM, RFL, AL, FCL

Supervision: GM, RFL, AL, FCL

Writing – original draft: ELdA, IS, FCL

Writing – review & editing: ELdA, IS, AFS, GM, RFL, AL, FCL

## Data and materials availability

All cell lines generated in this study are available upon request to FCL. Python and R scripts are available upon request. Processed data can be also visualized in our dedicated webtool: will be accessible upon publication of the work. All other data are available in the main text or the supplementary materials.

## Supplementary Materials

Figs. S1 to S7

Tables S3, S5 and S7 to S9

Auxiliary supplementary material: tables S1, S2, S4, S6 and S10.

## References and Notes

1. S. P. Jackson, J. Bartek, The DNA-damage response in human biology and disease. Nature 461, 1071–1078 (2009).

2. A. Ciccia, S. J. Elledge, The DNA Damage Response: Making It Safe to Play with Knives. Mol Cell 40, 179–204 (2010).

3. A. Tubbs, A. Nussenzweig, Endogenous DNA Damage as a Source of Genomic Instability in Cancer. Cell 168, 644–656 (2017).

4. A. Trenner, A. A. Sartori, Harnessing DNA Double-Strand Break Repair for Cancer Treatment. Front Oncol 9, 1–10 (2019).

5. J. Y. Wang, J. A. Doudna, CRISPR technology: A decade of genome editing is only the beginning. Science 379, eadd8643 (2023).

6. C. Xue, E. C. Greene, DNA Repair Pathway Choices in CRISPR-Cas9-Mediated Genome Editing. Trends in Genetics 37, 639–656 (2021).

7. M. van Overbeek, D. Capurso, M. M. Carter, M. S. Thompson, E. Frias, C. Russ, J. S. Reece-Hoyes, C. Nye, S. Gradia, B. Vidal, J. Zheng, G. R. Hoffman, C. K. Fuller, A. P. May, DNA Repair Profiling Reveals Nonrandom Outcomes at Cas9-Mediated Breaks. Mol Cell 63, 633–646 (2016).

8. M. W. Shen, M. Arbab, J. Y. Hsu, D. Worstell, S. J. Culbertson, O. Krabbe, C. A. Cassa, D. R. Liu, D. K. Gifford, R. I. Sherwood, Predictable and precise template-free CRISPR editing of pathogenic variants. Nature 563, 646–651 (2018).

9. A. Taheri-Ghahfarokhi, B. J. M. Taylor, R. Nitsch, A. Lundin, A. L. Cavallo, K. Madeyski-Bengtson, F. Karlsson, M. Clausen, R. Hicks, L. M. Mayr, M. Y. Bohlooly, M. Maresca, Decoding non-random mutational signatures at Cas9 targeted sites. Nucleic Acids Res 46, 8417–8434 (2018).

10. A. M. Chakrabarti, T. Henser-Brownhill, J. Monserrat, A. R. Poetsch, N. M. Luscombe, P. Scaffidi, Target-Specific Precision of CRISPR-Mediated Genome Editing. Mol Cell 73, 699–713.e6 (2019).

11. J. A. Hussmann, J. Ling, P. Ravisankar, J. Yan, A. Cirincione, A. Xu, D. Simpson, D. Yang, A. Bothmer, C. Cotta-Ramusino, J. S. Weissman, B. Adamson, Mapping the genetic landscape of DNA double-strand break repair. Cell 184, 5653–5669.e25 (2021).

12. J. Yan, P. Oyler-Castrillo, P. Ravisankar, C. C. Ward, S. Levesque, Y. Jing, D. Simpson, A. Zhao, H. Li, W. Yan, L. Goudy, R. Schmidt, S. C. Solley, L. A. Gilbert, M. M. Chan, D. E. Bauer, A. Marson, L. R. Parsons, B. Adamson, Improving prime editing with an endogenous small RNA-binding protein. Nature 628, 639– 647 (2024).

13. T. Hart, A. H. Y. Tong, K. Chan, J. Van Leeuwen, A. Seetharaman, M. Aregger, M. Chandrashekhar, N. Hustedt, S. Seth, A. Noonan, A. Habsid, O. Sizova, L. Nedyalkova, R. Climie, L. Tworzyanski, K. Lawson, M. A. Sartori, S. Alibeh, D. Tieu, S. Masud, P. Mero, A. Weiss, K. R. Brown, M. Usaj, M. Billmann, M. Rahman, M. Constanzo, C. L. Myers, B. J. Andrews, C. Boone, D. Durocher, J. Moffat, Evaluation and design of genome-wide CRISPR/SpCas9 knockout screens. G3: Genes, Genomes, Genetics 7, 2719–2727 (2017).

14. M. Olivieri, T. Cho, A. Álvarez-Quilón, K. Li, M. J. Schellenberg, M. Zimmermann, N. Hustedt, S. E. Rossi, S. Adam, H. Melo, A. M. Heijink, G. Sastre-Moreno, N. Moatti, R. K. Szilard, A. McEwan, A. K. Ling, A. Serrano-Benitez, T. Ubhi, S. Feng, J. Pawling, I. Delgado-Sainz, M. W. Ferguson, J. W. Dennis, G. W. Brown, F. Cortés-Ledesma, R. S. Williams, A. Martin, D. Xu, D. Durocher, A Genetic Map of the Response to DNA Damage in Human Cells. Cell 182, 481–496.e21 (2020).

15. X. Liu, T. Liu, Y. Shang, P. Dai, W. Zhang, B. J. Lee, M. Huang, D. Yang, Q. Wu, L. D. Liu, X. Zheng, B. O. Zhou, J. Dong, L. S. Yeap, J. Hu, T. Xiao, S. Zha, R. Casellas, X. S. Liu, F. L. Meng, ERCC6L2 promotes DNA orientation-specific recombination in mammalian cells. Cell Res 30, 732–744 (2020).

16. G. Sessa, B. Gómez-González, S. Silva, C. Pérez-Calero, R. Beaurepere, S. Barroso, S. Martineau, C. Martin, Å. Ehlén, J. S. Martínez, B. Lombard, D. Loew, S. Vagner, A. Aguilera, A. Carreira, BRCA2 promotes DNA-RNA hybrid resolution by DDX5 helicase at DNA breaks to facilitate their repair‡. EMBO J 40, 1–25 (2021).

17. J. S. Brown, N. Lukashchuk, M. Sczaniecka-Clift, S. Britton, C. le Sage, P. Calsou, P. Beli, Y. Galanty, S. P. Jackson, Neddylation Promotes Ubiquitylation and Release of Ku from DNA-Damage Sites. Cell Rep 11, 704–714 (2015).

18. I. Unk, I. Hajdú, A. Blastyák, L. Haracska, Role of yeast Rad5 and its human orthologs, HLTF and SHPRH in DNA damage tolerance. DNA Repair (Amst) 9, 257–267 (2010).

19. J. R. Lin, M. K. Zeman, J. Y. Chen, M. C. Yee, K. A. Cimprich, SHPRH and HLTF Act in a Damage-Specific Manner to Coordinate Different Forms of Postreplication Repair and Prevent Mutagenesis. Mol Cell 42, 237–249 (2011).

20. Y. J. Achar, D. Balogh, L. Haracska, Coordinated protein and DNA remodeling by human HLTF on stalled replication fork. Proc Natl Acad Sci U S A 108, 14073–14078 (2011).

21. A. C. Kile, D. A. Chavez, J. Bacal, S. Eldirany, D. M. Korzhnev, I. Bezsonova, B. F. Eichman, K. A. Cimprich, HLTF’s Ancient HIRAN Domain Binds 3’ DNA Ends to Drive Replication Fork Reversal. Mol Cell 58, 1090–1100 (2015).

22. G. Bai, C. Kermi, H. Stoy, C. J. Schiltz, J. Bacal, A. M. Zaino, M. K. Hadden, B. F. Eichman, M. Lopes, K. A. Cimprich, HLTF Promotes Fork Reversal, Limiting Replication Stress Resistance and Preventing Multiple Mechanisms of Unrestrained DNA Synthesis. Mol Cell 78, 1237–1251.e7 (2020).

23. I. Ahel, U. Rass, S. F. El-Khamisy, S. Katyal, P. M. Clements, P. J. McKinnon, K. W. Caldecott, S. C. West, The neurodegenerative disease protein aprataxin resolves abortive DNA ligation intermediates. Nature 443, 713–716 (2006).

24. A. W. Kijas, J. L. Harris, J. M. Harris, M. F. Lavin, Aprataxin forms a discrete branch in the HIT (Histidine Triad) superfamily of proteins with both DNA/RNA binding and nucleotide hydrolase activities. Journal of Biological Chemistry 281, 13939–13948 (2006).

25. U. Rass, I. Ahel, S. C. West, Actions of aprataxin in multiple DNA repair pathways. Journal of Biological Chemistry 282, 9469–9474 (2007).

26. P. Tumbale, J. S. Williams, M. J. Schellenberg, T. A. Kunkel, R. S. Williams, Aprataxin resolves adenylated RNA-DNA junctions to maintain genome integrity. Nature 506, 111–115 (2014).

27. L. Postow, H. Funabiki, An SCF complex containing Fbxl12 mediates DNA damage-induced Ku80 ubiquitylation. Cell Cycle 12, 587–595 (2013).

28. A. Brunner, Q. Li, S. Fisicaro, A. Kourtesakis, J. Viiliäinen, H. J. Johansson, V. Pandey, A. K. Mayank, J. Lehtiö, J. A. Wohlschlegel, C. Spruck, J. K. Rantala, L. M. Orre, O. Sangfelt, FBXL12 degrades FANCD2 to regulate replication recovery and promote cancer cell survival under conditions of replication stress. Mol Cell 83, 3720–3739.e8 (2023).

29. B. R. Lemos, A. C. Kaplan, J. E. Bae, A. E. Ferrazzoli, J. Kuo, R. P. Anand, D. P. Waterman, J. E. Haber, CRISPR/Cas9 cleavages in budding yeast reveal templated insertions and strand-specific insertion/deletion profiles. Proc Natl Acad Sci U S A 115, E2010–E2047 (2018).

30. M. M. Mehryar, X. Shi, J. Li, Q. Wu, DNA polymerases in precise and predictable CRISPR/Cas9-mediated chromosomal rearrangements. BMC Biol 21, 1–16 (2023).

31. A. Shibata, P. A. Jeggo, ATM’s role in the repair of DNA double-strand breaks. Genes (Basel) 12 (2021).

32. T. Ochi, A. N. Blackford, J. Coates, S. Jhujh, S. Mehmood, N. Tamura, J. Travers, Q. Wu, V. M. Draviam, C. V. Robinson, T. L. Blundell, S. P. Jackson, PAXX, a paralog of XRCC4 and XLF, interacts with Ku to promote DNA double-strand break repair. Science 347, 185–188 (2015).

33. V. Kumar, F. W. Alt, R. L. Frock, PAXX and XLF DNA repair factors are functionally redundant in joining DNA breaks in a G1-arrested progenitor B-cell line. Proc Natl Acad Sci U S A 113, 10619–10624 (2016).

34. C. Lescale, H. Lenden Hasse, A. N. Blackford, G. Balmus, J. J. Bianchi, W. Yu, L. Bacoccina, A. Jarade, C. Clouin, R. Sivapalan, B. Reina-San-Martin, S. P. Jackson, L. Deriano, Specific Roles of XRCC4 Paralogs PAXX and XLF during V(D)J Recombination. Cell Rep 16, 2967–2979 (2016).

35. P. J. Hung, B. R. Chen, R. George, C. Liberman, A. J. Morales, P. Colon-Ortiz, J. K. Tyler, B. P. Sleckman, A. L. Bredemeyer, Deficiency of XLF and PAXX prevents DNA double-strand break repair by non-homologous end joining in lymphocytes. Cell Cycle 16, 286–295 (2017).

36. M. Seif-El-Dahan, A. Kefala-Stavridi, P. Frit, S. W. Hardwick, D. Y. Chirgadze, T. M. De Oliviera, S. Britton, N. Barboule, M. Bossaert, A. P. Pandurangan, K. Meek, T. L. Blundell, V. Ropars, P. Calsou, J. B. Charbonnier, A. K. Chaplin, PAXX binding to the NHEJ machinery explains functional redundancy with XLF. Sci Adv 9, 1–14 (2023).

37. S. Chen, A. Vogt, L. Lee, T. Naila, R. McKeown, A. E. Tomkinson, S. P. Lees-Miller, Y. He, Cryo-EM visualization of DNA-PKcs structural intermediates in NHEJ. Sci Adv 9, 1–10 (2023).

38. J. Zhang, Q. Zhang, VHL and hypoxia signaling: Beyond HIF in cancer. Biomedicines 6, 1–13 (2018).

39. S. M. Noordermeer, S. Adam, D. Setiaputra, M. Barazas, S. J. Pettitt, A. K. Ling, M. Olivieri, A. Álvarez-Quilón, N. Moatti, M. Zimmermann, S. Annunziato, D. B. Krastev, F. Song, I. Brandsma, J. Frankum, R. Brough, A. Sherker, S. Landry, R. K. Szilard, M. M. Munro, A. McEwan, T. G. de Rugy, Z. Y. Lin, T. Hart, J. Moffat, A. C. Gingras, A. Martin, H. van Attikum, J. Jonkers, C. J. Lord, S. Rottenberg, D. Durocher, The shieldin complex mediates 53BP1-dependent DNA repair. Nature 560, 117–121 (2018).

40. H. Dev, T. W. W. Chiang, C. Lescale, I. de Krijger, A. G. Martin, D. Pilger, J. Coates, M. Sczaniecka-Clift, W. Wei, M. Ostermaier, M. Herzog, J. Lam, A. Shea, M. Demir, Q. Wu, F. Yang, B. Fu, Z. Lai, G. Balmus, R. Belotserkovskaya, V. Serra, M. J. O’Connor, A. Bruna, P. Beli, L. Pellegrini, C. Caldas, L. Deriano, J. J. L. Jacobs, Y. Galanty, S. P. Jackson, Shieldin complex promotes DNA end-joining and counters homologous recombination in BRCA1-null cells. Nat Cell Biol 20, 954–965 (2018).

41. R. Gupta, K. Somyajit, T. Narita, E. Maskey, A. Stanlie, M. Kremer, D. Typas, M. Lammers, N. Mailand, A. Nussenzweig, J. Lukas, C. Choudhary, DNA Repair Network Analysis Reveals Shieldin as a Key Regulator of NHEJ and PARP Inhibitor Sensitivity. Cell 173, 972–988.e23 (2018).

42. H. Ghezraoui, C. Oliveira, J. R. Becker, K. Bilham, D. Moralli, C. Anzilotti, R. Fischer, M. Deobagkar-Lele, M. Sanchiz-Calvo, E. Fueyo-Marcos, S. Bonham, B. M. Kessler, S. Rottenberg, R. J. Cornall, C. M. Green, J. R. Chapman, 53BP1 cooperation with the REV7–shieldin complex underpins DNA structure-specific NHEJ. Nature 560, 122–127 (2018).

43. S. Findlay, J. Heath, V. M. Luo, A. Malina, T. Morin, Y. Coulombe, B. Djerir, Z. Li, A. Samiei, E. Simo-Cheyou, M. Karam, H. Bagci, D. Rahat, D. Grapton, E. G. Lavoie, C. Dove, H. Khaled, H. Kuasne, K. K. Mann, K. O. Klein, C. M. Greenwood, Y. Tabach, M. Park, J. Côté, J. Masson, A. Maréchal, A. Orthwein, SHLD 2/ FAM 35A co-operates with REV 7 to coordinate DNA double-strand break repair pathway choice. EMBO J 37, 1–20 (2018).

44. J. Tomida, K. Takata, S. Bhetawal, M. D. Person, H. Chao, D. G. Tang, R. D. Wood, FAM 35A associates with REV 7 and modulates DNA damage responses of normal and BRCA 1-defective cells. EMBO J 37, 1–14 (2018).

45. K. M. Christopher J. Buehl, Noah J. Goff1, Steven W. Hardwick, Martin Gellert, Tom L. Blundell, Wei Yang, Amanda K. Chaplin, Two distinct long-range synaptic complexes promote different aspects of end-processing prior to repair of DNA breaks by non-homologous end joining. Mol Cell 83, 698–714 (2023).

46. S. Chen, L. Lee, T. Naila, S. Fishbain, A. Wang, A. E. Tomkinson, S. P. Lees-Miller, Y. He, Structural basis of long-range to short-range synaptic transition in NHEJ. Nature 593, 294–298 (2021).

47. A. K. Chaplin, S. W. Hardwick, S. Liang, A. Kefala Stavridi, A. Hnizda, L. R. Cooper, T. M. De Oliveira, D. Y. Chirgadze, T. L. Blundell, Dimers of DNA-PK create a stage for DNA double-strand break repair. Nat Struct Mol Biol 28, 13–19 (2021).

48. A. K. Chaplin, S. W. Hardwick, A. K. Stavridi, C. J. Buehl, N. J. Goff, V. Ropars, S. Liang, T. M. De Oliveira, D. Y. Chirgadze, K. Meek, J. B. Charbonnier, T. L. Blundell, Cryo-EM of NHEJ supercomplexes provides insights into DNA repair. Mol Cell 81, 3400–3409.e3 (2021).

49. A. Fradet-Turcotte, M. D. Canny, C. Escribano-Díaz, A. Orthwein, C. C. Y. Leung, H. Huang, M. C. Landry, J. Kitevski-Leblanc, S. M. Noordermeer, F. Sicheri, D. Durocher, 53BP1 is a reader of the DNA-damage-induced H2A Lys 15 ubiquitin mark. Nature 499, 50–54 (2013).

50. R. S. Zou, A. Marin-Gonzalez, Y. Liu, H. B. Liu, L. Shen, R. K. Dveirin, J. X. J. Luo, R. Kalhor, T. Ha, Massively parallel genomic perturbations with multi-target CRISPR interrogates Cas9 activity and DNA repair at endogenous sites. Nat Cell Biol 24, 1433–1444 (2022).

51. J. Shou, J. Li, Y. Liu, Q. Wu, Precise and Predictable CRISPR Chromosomal Rearrangements Reveal Principles of Cas9-Mediated Nucleotide Insertion. Mol Cell 71, 498–509.e4 (2018).

52. X. Shi, J. Shou, M. M. Mehryar, J. Li, L. Wang, M. Zhang, H. Huang, X. Sun, Q. Wu, Cas9 has no exonuclease activity resulting in staggered cleavage with overhangs and predictable di- and tri-nucleotide CRISPR insertions without template donor. Cell Discov 5, 4–7 (2019).

53. I. Unk, I. Hajdú, K. Fátyol, J. Hurwitz, J. H. Yoon, L. Prakash, S. Prakash, L. Haracska, Human HLTF functions as a ubiquitin ligase for proliferating cell nuclear antigen polyubiquitination. Proc Natl Acad Sci U S A 105, 3768–3773 (2008).

54. Y. Masuda, M. Suzuki, H. Kawai, A. Hishiki, H. Hashimoto, C. Masutani, T. Hishida, F. Suzuki, K. Kamiya, En bloc transfer of polyubiquitin chains to PCNA in vitro is mediated by two different human E2-E3 pairs. Nucleic Acids Res 40, 10394–10407 (2012).

55. A. Hishiki, K. Hara, Y. Ikegaya, H. Yokoyama, T. Shimizu, M. Sato, H. Hashimoto, Structure of a novel DNA-binding domain of Helicase-like Transcription Factor (HLTF) and its functional implication in DNA damage tolerance. Journal of Biological Chemistry 290, 13215–13223 (2015).

56. D. A. Chavez, B. H. Greer, B. F. Eichman, The HIRAN domain of helicase-like transcription factor positions the DNA translocase motor to drive efficient DNA fork regression. Journal of Biological Chemistry 293, 8484–8494 (2018).

57. J. J. Hsieh, M. P. Purdue, S. Signoretti, C. Swanton, L. Albiges, M. Schmidinger, D. Y. Heng, J. Larkin, V. Ficarra, Renal cell carcinoma. Nat Rev Dis Primers 3, 1–19 (2017).

58. S. Turajlic, K. Litchfield, H. Xu, R. Rosenthal, N. McGranahan, J. L. Reading, Y. N. S. Wong, A. Rowan, N. Kanu, M. Al Bakir, T. Chambers, R. Salgado, P. Savas, S. Loi, N. J. Birkbak, L. Sansregret, M. Gore, J. Larkin, S. A. Quezada, C. Swanton, Insertion-and-deletion-derived tumour-specific neoantigens and the immunogenic phenotype: a pan-cancer analysis. Lancet Oncol 18, 1009–1021 (2017).

59. L. B. Alexandrov, J. Kim, N. J. Haradhvala, M. N. Huang, A. W. Tian Ng, Y. Wu, A. Boot, K. R. Covington, D. A. Gordenin, E. N. Bergstrom, S. M. A. Islam, N. Lopez-Bigas, L. J. Klimczak, J. R. McPherson, S. Morganella, R. Sabarinathan, D. A. Wheeler, V. Mustonen, P. Boutros, K. Chan, A. Fujimoto, G. Getz, M. N. Huang, M. Kazanov, M. Lawrence, I. Martincorena, S. Morganella, H. Nakagawa, P. Polak, S. Prokopec, S. A. Roberts, S. G. Rozen, N. Saini, T. Shibata, Y. Shiraishi, M. R. Stratton, B. T. Teh, I. Vázquez-García, F. Yousif, W. Yu, The repertoire of mutational signatures in human cancer. Nature 578, 94–101 (2020).

60. P. J. Chen, J. A. Hussmann, J. Yan, F. Knipping, P. Ravisankar, P. F. Chen, C. Chen, J. W. Nelson, G. A. Newby, M. Sahin, M. J. Osborn, J. S. Weissman, B. Adamson, D. R. Liu, Enhanced prime editing systems by manipulating cellular determinants of editing outcomes. Cell 184, 5635–5652.e29 (2021).

61. L. W. Koblan, M. Arbab, M. W. Shen, J. A. Hussmann, A. V. Anzalone, J. L. Doman, G. A. Newby, D. Yang, B. Mok, J. M. Replogle, A. Xu, T. A. Sisley, J. S. Weissman, B. Adamson, D. R. Liu, Efficient C•G-to-G•C base editors developed using CRISPRi screens, target-library analysis, and machine learning. Nat Biotechnol 39, 1414–1425 (2021).

62. G. Balmus, A. C. Barros, P. W. G. Wijnhoven, C. Lescale, H. L. Hasse, K. Boroviak, C. le Sage, B. Doe, A. O. Speak, A. Galli, M. Jacobsen, L. Deriano, D. J. Adams, A. N. Blackford, S. P. Jackson, Synthetic lethality between PAXX and XLF in mammalian development. Genes Dev 30, 2152–2157 (2016).

63. G. M. C. Longo, S. Sayols, A. G. Kotini, S. Heinen, M. M. Möckel, P. Beli, V. Roukos, Linking CRISPR– Cas9 Double-Strand Break Profiles to Gene Editing Precision with BreakTag. Nat Biotechnol (2024). Online ahead of print

64. S. S. Chen, A. A. Davies, D. Sagan, H. D. Ulrich, The RING finger ATPase Rad5p of Saccharomyces cerevisiae contributes to DNA double-strand break repair in a ubiquitin-independent manner. Nucleic Acids Res 33, 5878–5886 (2005).

65. F. Li, A. Zafar, L. Luo, A. M. Denning, J. Gu, A. Bennett, F. Yuan, Y. Zhang, R-Loops in Genome Instability and Cancer. Cancers (Basel) 15 (2023).

66. J. Domingo-Prim, F. Bonath, N. Visa, RNA at DNA Double-Strand Breaks: The Challenge of Dealing with DNA:RNA Hybrids. BioEssays 42, 1–7 (2020).

67. J. H. Seol, E. Y. Shim, S. E. Lee, Microhomology-mediated end joining: Good, bad and ugly. Mutation 809, 81–87 (2018).

68. H. Fleury, M. K. MacEachern, C. M. Stiefel, R. Anand, C. Sempeck, B. Nebenfuehr, K. Maurer-Alcalá, K. Ball, B. Proctor, O. Belan, E. Taylor, R. Ortega, B. Dodd, L. Weatherly, D. Dansoko, J. W. Leung, S. J. Boulton, N. Arnoult, The APE2 nuclease is essential for DNA double-strand break repair by microhomology-mediated end joining. Mol Cell 83, 1429–1445.e8 (2023).

69. A. Brambati, O. Sacco, S. Porcella, J. Heyza, M. Kareh, J. C. Schmidt, A. Sfeir, RHINO directs MMEJ to repair DNA breaks in mitosis. Science (1979) 381, 653–660 (2023).

70. A. V. Nimonkar, J. Genschel, E. Kinoshita, P. Polaczek, J. L. Campbell, C. Wyman, P. Modrich, S. C. Kowalczykowski, BLM-DNA2-RPA-MRN and EXO1-BLM-RPA-MRN constitute two DNA end resection machineries for human DNA break repair. Genes Dev 25, 350–362 (2011).

71. S. Nath, G. Nagaraju, FANCJ Helicase Promotes DNA End Resection by Facilitating CtIP Recruitment to DNA Double-Strand Breaks (2020; 10.1371/journal.pgen.1008701)vol. 16.

72. M. Y. Cai, C. E. Dunn, W. Chen, B. S. Kochupurakkal, H. Nguyen, L. A. Moreau, G. I. Shapiro, K. Parmar, D. Kozono, A. D. D’Andrea, Cooperation of the ATM and Fanconi Anemia/BRCA Pathways in Double-Strand Break End Resection. Cell Rep 30, 2402–2415.e5 (2020).

73. M. M. Soniat, G. Nguyen, H. C. Kuo, I. J. Finkelstein, The MRN complex and topoisomerase IIIa–RMI1/2 synchronize DNA resection motor proteins. Journal of Biological Chemistry 299, 102802 (2023).

74. X. Cheng, V. R. Cote, J. Côte, NuA4 and SAGA acetyltransferase complexes cooperate for repair of DNA breaks by homologous recombination. PLoS Genet 17, 1–19 (2021).

75. T. Clouaire, V. Rocher, A. Lashgari, C. Arnould, M. Aguirrebengoa, A. Biernacka, M. Skrzypczak, F. Aymard, B. Fongang, N. Dojer, J. S. Iacovoni, M. Rowicka, K. Ginalski, J. Côté, G. Legube, Comprehensive Mapping of Histone Modifications at DNA Double-Strand Breaks Deciphers Repair Pathway Chromatin Signatures. Mol Cell 72, 250–262.e6 (2018).

76. A. Schrempf, J. Slyskova, J. I. Loizou, Targeting the DNA Repair Enzyme Polymerase θ in Cancer Therapy. Trends Cancer 7, 98–111 (2021).

77. M. Tang, E. Bolderson, K. J. O’Byrne, D. J. Richard, Tumor Hypoxia Drives Genomic Instability. Front Cell Dev Biol 9 (2021).

78. J. P. Concordet, M. Haeussler, CRISPOR: Intuitive guide selection for CRISPR/Cas9 genome editing experiments and screens. Nucleic Acids Res 46, W242–W245 (2018).

79. E. K. Brinkman, T. Chen, M. Amendola, B. Van Steensel, Easy quantitative assessment of genome editing by sequence trace decomposition. Nucleic Acids Res 42, e168 (2014).

80. M. Olivieri, Genome-scale chemogenomic CRISPR screens in human cells using the TKOv3 library. STAR Protoc 2, 100321 (2021).

81. J. García-Nafría, J. F. Watson, I. H. Greger, IVA cloning: A single-tube universal cloning system exploiting bacterial In Vivo Assembly. Sci Rep 6, 1–12 (2016).

82. W. Feng, D. A. Simpson, J. E. Cho, J. Carvajal-Garcia, C. M. Smith, K. M. Headley, N. Hathaway, D. A. Ramsden, G. P. Gupta, Marker-free quantification of repair pathway utilization at Cas9-induced double-strand breaks. Nucleic Acids Res 49, 5095–5105 (2021).

83. V. Bhandari, C. H. Li, R. G. Bristow, P. C. Boutros, PCAWG Consortium, Divergent mutational processes distinguish hypoxic and normoxic tumours. Nat Commun 11, 1–10 (2020).

84. F. M. Buffa, A. L. Harris, C. M. West, C. J. Miller, Large meta-analysis of multiple cancers reveals a common, compact and highly prognostic hypoxia metagene. Br J Cancer 102, 428–435 (2010).

