## Supplementary figures and tables for "A comprehensive genetic catalog of human double-strand break repair"

**The PDF file includes:**

Figs. S1 to S7
Tables S3, S5, S7 to S9

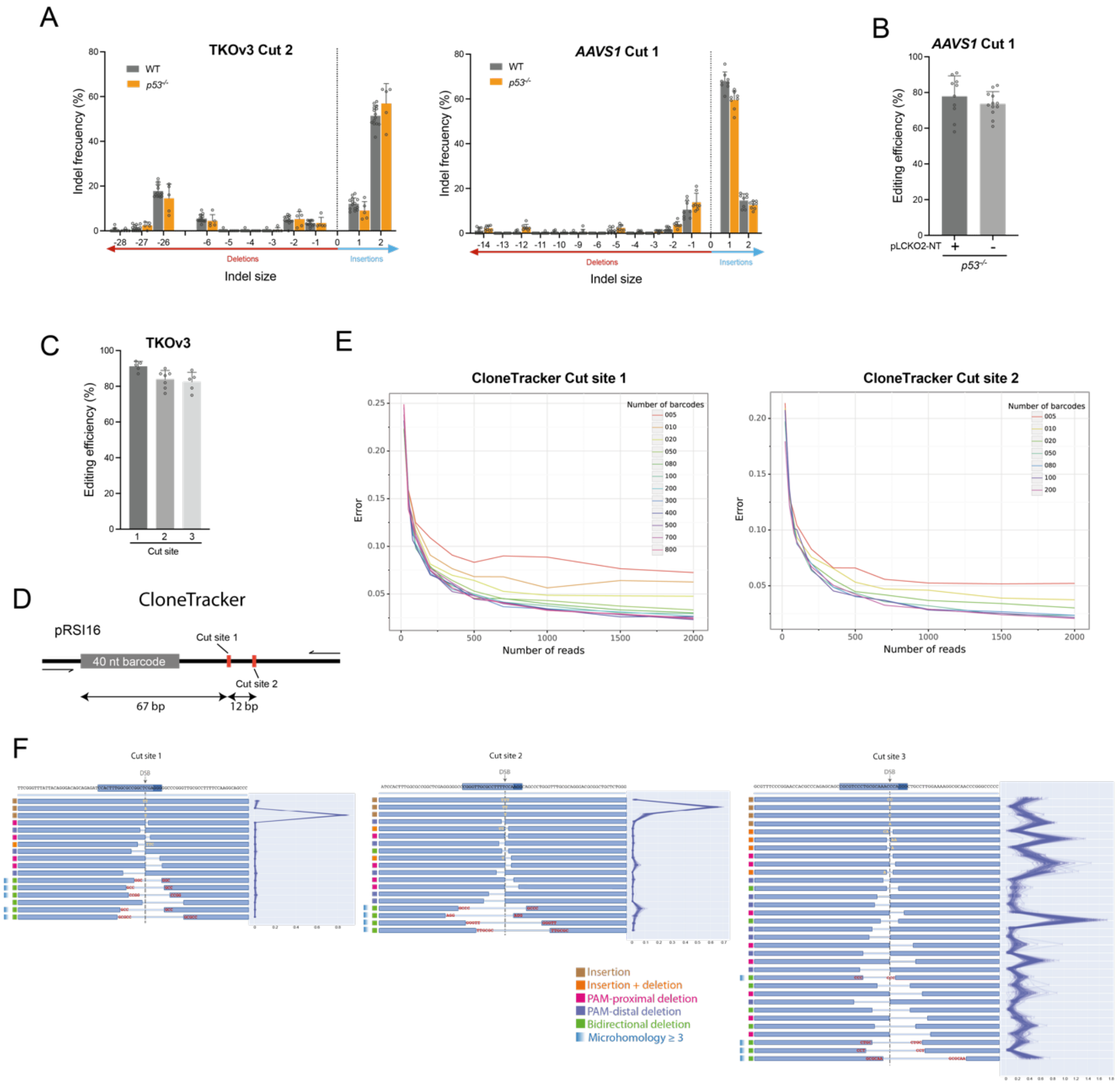

A
TKOv3 Cut 2
AAVS1 Cut 1
WT
p53-/-
Indel frecuency (%)
Deletions
Insertions
Indel size
B
AAVS1 Cut 1
Editing efficiency (%)
pLCKO2-NT
p53-/-
C
TKOv3
Editing efficiency (%)
Cut site
D
CloneTracker
pRSI16
40 nt barcode
Cut site 1
Cut site 2
67 bp
12 bp
E
CloneTracker Cut site 1
CloneTracker Cut site 2
Number of barcodes
Error
Number of reads
F
Cut site 1
Cut site 2
Cut site 3
DSB
Insertion
Insertion + deletion
PAM-proximal deletion
PAM-distal deletion
Bidirectional deletion
Microhomology ≥ 3

**Fig. S1. Determination of REPAIRome screen settings, experimental representation and depth.**

**(A)** Distribution of most frequent indels produced at TKOv3 Cut site 2 (left) and at *AAVS1* Cut site 1 (right) in RPE1-*Cas9* TKOv3-NT and RPE1-*Cas9* cells, respectively, either deficient or proficient. Experiments carried out as in Fig. 1C. n = 14 for *WT* TKOv3-NT, 5 for *p53*$^{-/-}$ TKOv3-NT and 8 for *WT* and *p53*$^{-/-}$ at the *AAVS1* Cut site 1. Data show average of indel frequency, with error bars representing ±SD.

**(B)** Editing efficiency (measured as the percentage of sequencing reads containing indels at the cut site from the total of sequenced events) estimated at the *AAVS1* Cut 1 3 days after gRNA transfection, in *p53*$^{-/-}$ cells with and without the pLCKO2-NT lentiviral plasmid integrated in the genome, i.e., expressing or not a non-targeting sgRNA. n = 10 for *p53*$^{+/+}$ and 12 for *p53*$^{-/-}$. Error bars represent ±SD.

**(C)** Editing efficiency measured 3 days after transfection with gRNAs targeting each of the three cut sites analysed in the screen in the RPE1-*Cas9* TKOv3-NT cell line. n = 5, 7 and 5 for TKOv3 cut sites 1, 2 and 3, respectively. Error bars represent ±SD.

**(D)** Schematic representation of the CloneTracker barcode library lentivector cassette used for determining the screen experimental depth. The location of the cut sites tested and primers used to generate the Illumina library are indicated.

**(E)** Variability (error) in indel frequency distribution between virtual replicates when randomly selecting increasing number of independent barcodes and total repair events (reads) for the two cut sites shown in Fig.S1D.

**(F)** Diagrams depicting the most frequent indels for each cut analysed in the screen (left) and their relative frequency for the control sgRNAs (right). Protospacer and PAM sequences are highlighted in blue on the top. The expected cut site for each sgRNA is indicated by a vertical dashed line. Inserted nucleotides are shown in yellow and microhomology regions (≥ 3 nucleotides) flanking a deletion are shown in red. The category of each indel is indicated by coloured squares on the left side of the diagram.

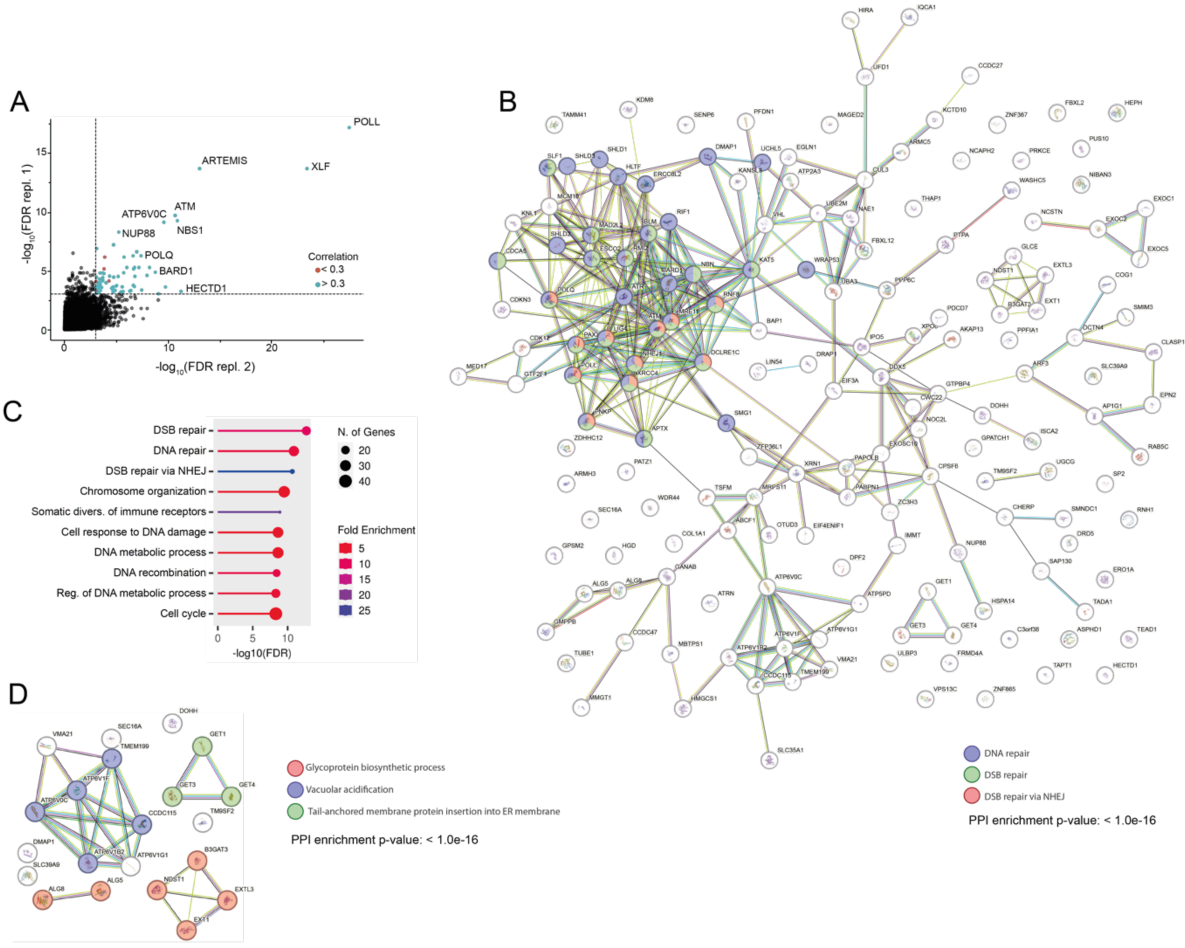
A
POLL
ARTEMIS
XLF
ATM
ATP6V0C
NBS1
NUP88
POLQ
BARD1
HECTD1
Correlation
< 0.3
> 0.3
-log10(FDR repl. 1)
-log10(FDR repl. 2)
B
DNA repair
DSB repair
DSB repair via NHEJ
PPI enrichment p-value: < 1.0e-16
C
DSB repair
DNA repair
DSB repair via NHEJ
Chromosome organization
Somatic divers. of immune receptors
Cell response to DNA damage
DNA metabolic process
DNA recombination
Reg. of DNA metabolic process
Cell cycle
N. of Genes
20
30
40
Fold Enrichment
5
10
15
20
25
-log10(FDR)
D
Glycoprotein biosynthetic process
Vacuolar acidification
Tail-anchored membrane protein insertion into ER membrane
PPI enrichment p-value: < 1.0e-16

**Fig. S2. Screen data processing and analysis.**

**(A)** Example of gene filtering for TKOv3 Cut site 2 using FDR < 0.01 and PCC between replicates > 0.3 as eligibility criteria.

**(B)** STRING analysis of functional interactions between the selected genes with the highest impact in the repair pattern when knocked out, i.e., FDR < 0.01 and PCC between replicates > 0.35, for at least one cut, and an overall distance to controls > 5 (Table S2). Proteins previously associated with GO terms "DNA repair", "DSB repair" and "DSB repair via NHEJ" are coloured as indicated.

**(C)** Gene ontology enrichment of genes selected in Table S2. Analysis performed with ShinyGO (version 0.77).

**(D)** STRING analysis of functional interactions between genes included in the low editing-efficiency cluster.

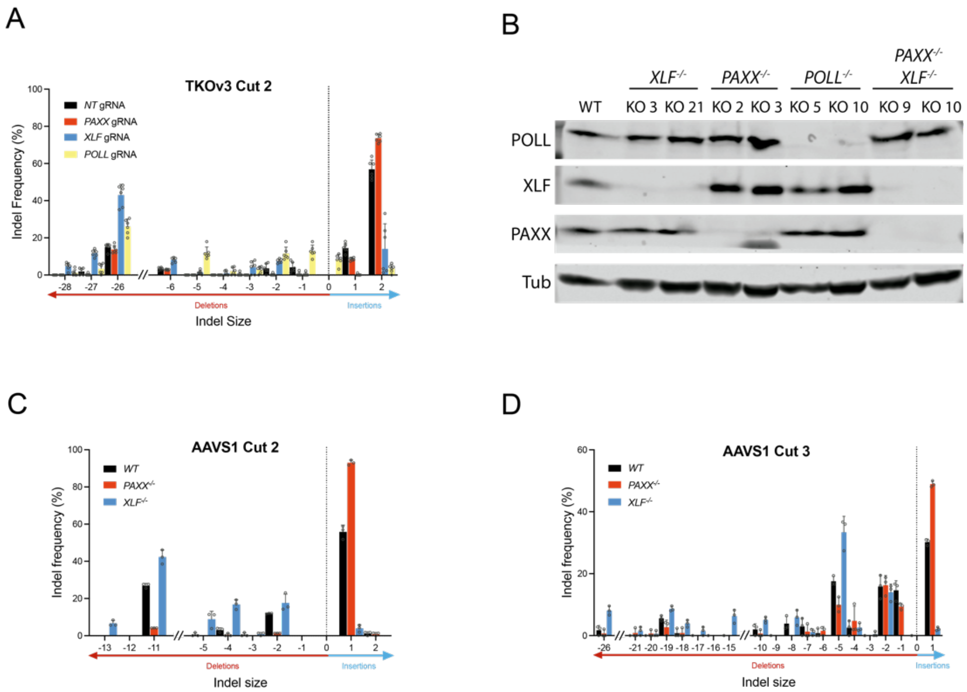
A
TKOv3 Cut 2
NT gRNA
PAXX gRNA
XLF gRNA
POLL gRNA
Indel Frequency (%)
Deletions
Insertions
Indel Size
B
XLF-/-
PAXX-/-
POLL-/-
PAXX-/- XLF-/-
WT
KO 3
KO 21
KO 2
KO 3
KO 5
KO 10
KO 9
KO 10
POLL
XLF
PAXX
Tub
C
AAVS1 Cut 2
WT
PAXX-/-
XLF-/-
Indel frequency (%)
Deletions
Insertions
Indel size
D
AAVS1 Cut 3
WT
PAXX-/-
XLF-/-
Indel frequency (%)
Deletions
Insertions
Indel size

**Fig. S3. Validation of the PAXX/XLF phenotype at different Cas9 target sites.**

**(A)** Effects of indicated *de-novo* designed gRNAs transfection on the repair profile at TKOv3 Cut site 2 in RPE1-*Cas9* TKOv3-NT cells, shown as in (Fig. 1C). Cells are transfected with the indicated gRNAs to create the desired polyclonal knockout populations and kept cycling for 12 days, to allow for protein clearance. Then, cells are transfected with the TKOv3 Cut 2-targeting gRNA and genomic DNA is purified three days later. Both the TKOv3 Cut2 repair pattern and the percentage of knockout in the polyclonal population (Table S3) are determined by Sanger sequencing and Sanger trace deconvolution software (Synthego-ICE). Data show average values normalized to total editing efficiency for 2 *de-novo* gRNA per gene and 3 independent replicates of each gRNA for each gene. Error bars show ±SD

**(B)** Immunoblotting for *PAXX, XLF* and *POLL* to verify the indicated knockout clones. Tubulin was used as a loading control.

**(C)** Effect of PAXX and XLF on the repair profile at two endogenous Cas9 targets located in the *AAVS1* region. Assay performed as in Fig. 1C. n = 3, error bars show ±SD.

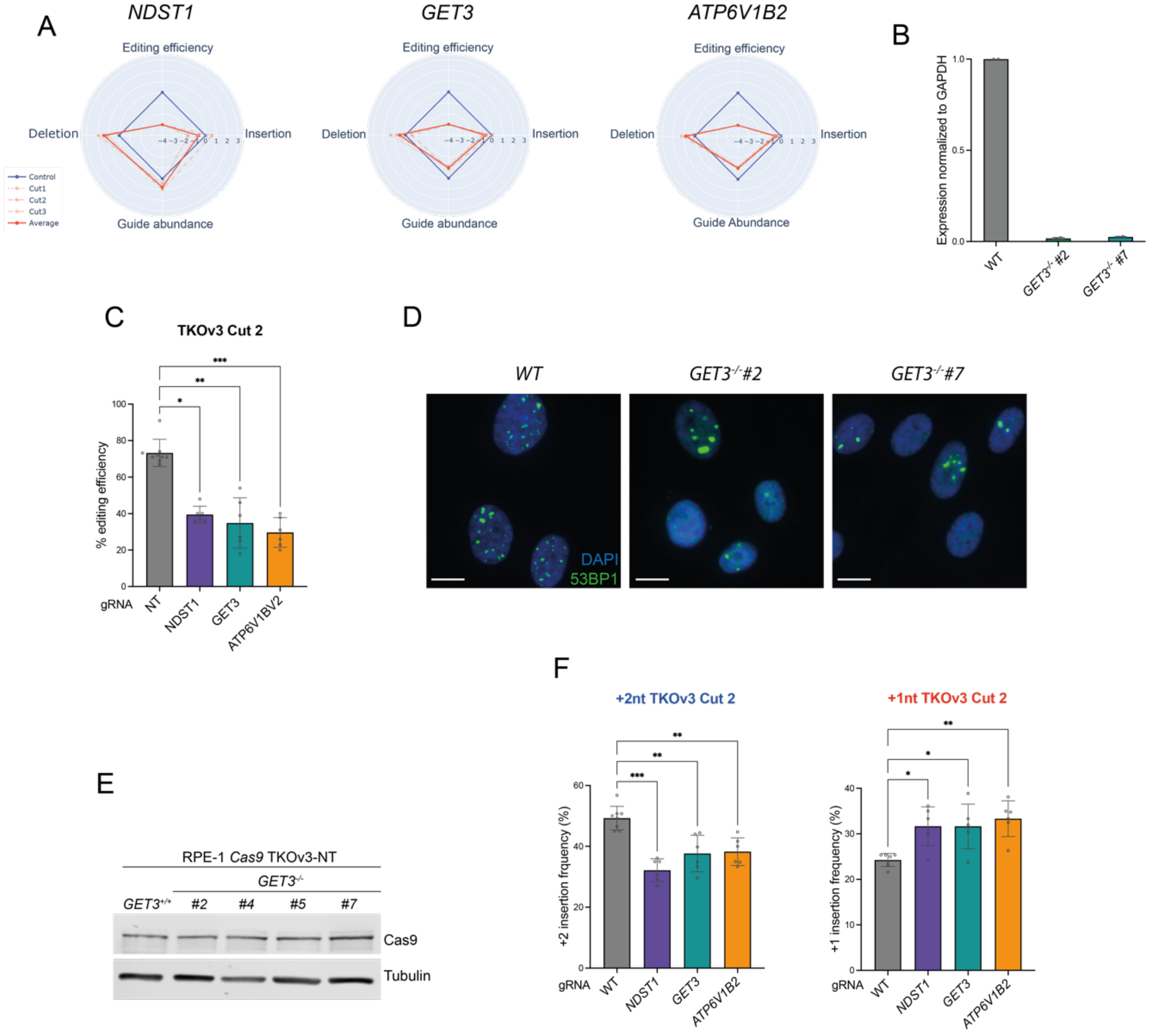

A
NDST1
GET3
ATP6V1B2
Editing efficiency
Insertion
Deletion
Guide abundance
Guide Abundance
Control
Cut1
Cut2
Cut3
Average
B
Expression normalized to GAPDH
WT
GET3-/- #2
GET3-/- #7
C
TKOv3 Cut 2
% editing efficiency
gRNA
NT
NDST1
GET3
ATP6V1BV2
D
WT
GET3-/-#2
GET3-/-#7
DAPI
53BP1
E
RPE-1 Cas9 TKOv3-NT
GET3-/-
GET3+/+
#2
#4
#5
#7
Cas9
Tubulin
F
+2nt TKOv3 Cut 2
+1nt TKOv3 Cut 2
+2 insertion frquency (%)
+1 insertion frequency (%)
gRNA
WT
NDST1
GET3
ATP6V1B2

**Fig. S4. Validation of decreased editing efficiency gene cluster.**

**(A)** Radar plots displaying the change of the specified parameters in the indicated genetic backgrounds. Values show the normalized frequency change of the indicated sgRNAs (red lines) relative to controls (blue line). The continuous red line shows the average change of the three cut sites combined, while the dashed lines show the changes for each independent cut site.

**(B)** Quantification by RT-qPCR of *GET3* mRNA levels in the *GET3*$^{-/-}$ candidate clones relative to wild type cells. Amplification levels were normalized to those of GAPDH. n = 2, error bars show ±SD.

**(C)** Effects of indicated *de-novo* designed gRNAs transfection on the editing efficiency at TKOv3 Cut site 2 in RPE1-*Cas9* TKOv3-NT cells. Assay performed as in Fig. S3A, but using half of the usual TKOv3 Cut site 2 gRNA concentration (6 nM), at which the reduction in editing efficiency becomes more evident. Data show average values normalized to total editing efficiency for 2 different guides per gene and 3 independent replicates of each guide. Error bars show ±SD. Statistical analysis was performed using One-way ANOVA.

**(D)** Representative immunofluorescence images of experiment corresponding to Fig. 3C, showing 53BP1 foci in RPE1-*Cas9* TKOv3-NT cells either deficient or proficient for *GET3.* Cells are transfected with the multisite gRNA (1.5 nM) and fixed 12h later. Scale bar represents 10 µm.

**(E)** Immunoblotting for Cas9 in the RPE1-Cas9 TKOv3-NT cell line and its *GET3*$^{-/-}$ derivative clones. Tubulin was used as a loading control.

**(F)** Effects of indicated *de-novo* designed gRNAs transfection on the frequency of +2 (left) and +1 (right) nucleotide insertions at TKOv3 Cut site 2 in RPE1-*Cas9* TKOv3-NT cells. Assay performed as in Fig. S3A. Data show average values normalized to total editing efficiency for 2 different guides per gene and 3 independent replicates of each guide. Error bars show ±SD. Statistical analysis was performed using One-way ANOVA.

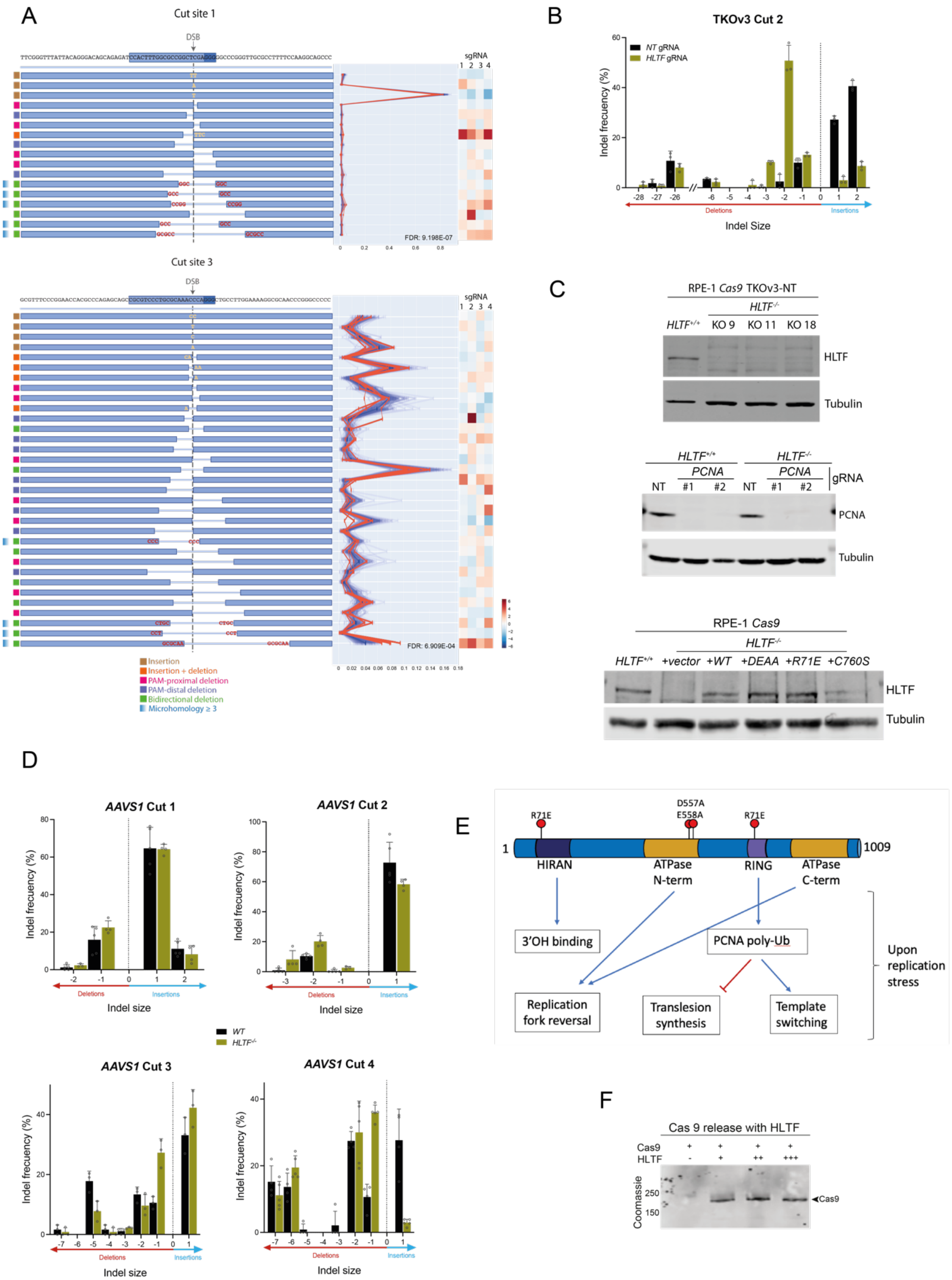
A
Cut site 1
DSB
Cut site 3
sgRNA
Insertion
Insertion + deletion
PAM-proximal deletion
PAM-distal deletion
Bidirectional deletion
Microhomology ≥ 3
B
TKOv3 Cut 2
NT gRNA
HLTF gRNA
Indel frecuency (%)
Deletions
Insertions
Indel Size
C
RPE-1 Cas9 TKOv3-NT
HLTF-/-
HLTF+/+
KO 9
KO 11
KO 18
HLTF
Tubulin
PCNA
gRNA
NT
#1
#2
RPE-1 Cas9
+vector
+WT
+DEAA
+R71E
+C760S
D
AAVS1 Cut 1
AAVS1 Cut 2
AAVS1 Cut 3
AAVS1 Cut 4
Indel size
WT
HLTF-/-
E
R71E
D557A
E558A
1
1009
HIRAN
ATPase N-term
RING
ATPase C-term
3'OH binding
PCNA poly-Ub
Replication fork reversal
Translesion synthesis
Template switching
Upon replication stress
F
Cas 9 release with HLTF
Cas9
HLTF
Coomassie
250
150
Cas9

**Fig. S5. HLTF effect on the repair of Cas9-induced DSBs.**

**(A)** Summary of results obtained in the screen for HLTF, as in Fig. 1D. Data for TKOv3 Cut site 1 and 3 is displayed.

**(B)** Effects of indicated *de-novo* designed gRNAs transfection on indels distribution produced at TKOv3 Cut site 2 in RPE1-*Cas9* TKOv3-NT cells. Assay carried out as in Fig. S3A. n = 3, error bars show ±SD.

**(C)** Immunoblotting for HLTF to verify knockout clones in the RPE1-*Cas9* TKOv3-NT background (top); for PCNA to corroborate its depletion in experiments corresponding to Fig. 5C (centre); and for HLTF to check its expression induction in the RPE1-*Cas9* *HLTF*$^{-/-}$ cells complemented with the different mutant versions of *HLTF* (bottom). Tubulin was used as a loading control.

**(D)** Repair profiles obtained at four endogenous Cas9 target sites located in the *AAVS1* region for the RPE1-*Cas9* wild type and *HLTF* knockout cell lines. Assay performed as in Fig. 1C. n = 4 for cut sites 1 and 2, n = 3 for cut site 3; and n = 5 for cut site 4. Error bars show ±SD.

**(E)** Schematic representation of HLTF architecture with its functional domains and their most relevant functions. The different mutations tested in this work are indicated with red pins.

**(F)** In vitro removal of post-cleavage Cas9 RNP by HLTF (see Fig. 5 and Methods section). Briefly, immobilized TKOv3 cut site 2 sequence is digested with its targeting Cas9 RNP. Then, post-cleavage DNA-bound Cas9 RNP is incubated in the presence of increasing amounts of HLTF purified from insect cells. Finally, Cas9 in the flow-through material from this mix is analysed by SDS-PAGE gel electrophoresis and western blotting.

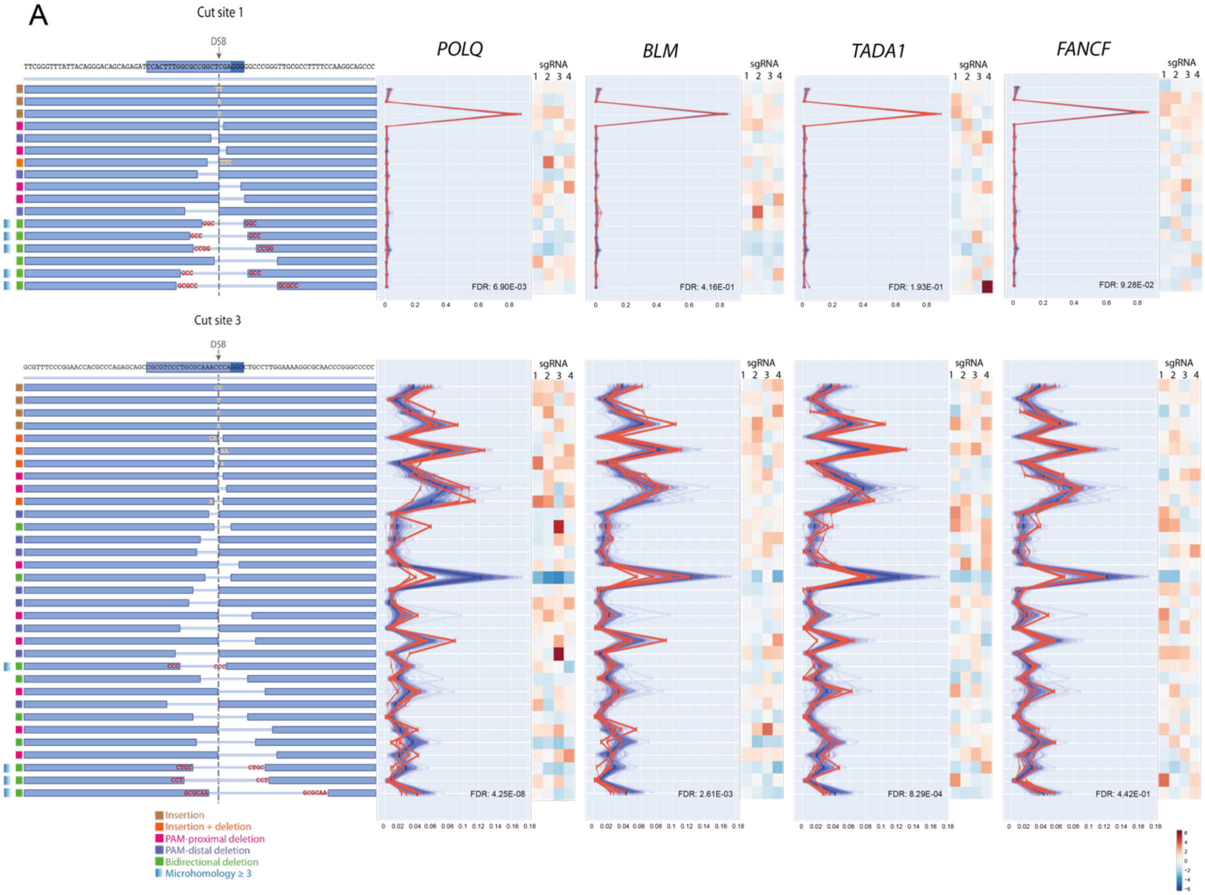

A
Cut site 1
DSB
POLQ
BLM
TADA1
FANCF
sgRNA
1 2 3 4
FDR: 6.90E-03
FDR: 4.16E-01
FDR: 1.93E-01
FDR: 9.28E-02
Cut site 3
FDR: 4.25E-08
FDR: 2.61E-03
FDR: 8.29E-04
FDR: 4.42E-01
Insertion
Insertion + deletion
PAM-proximal deletion
PAM-distal deletion
Bidirectional deletion
Microhomology ≥ 3

**Fig. S6. Screen results for representative genes of the *POLQ* correlation network.**

**(A)** Summary of results obtained in the screen for selected genes (*POLQ, BLM, TADA1* and FANCF) from the *POLQ* correlation network (shown in fig. 6B). Data for TKOv3 Cut site 1 and 3 is displayed.

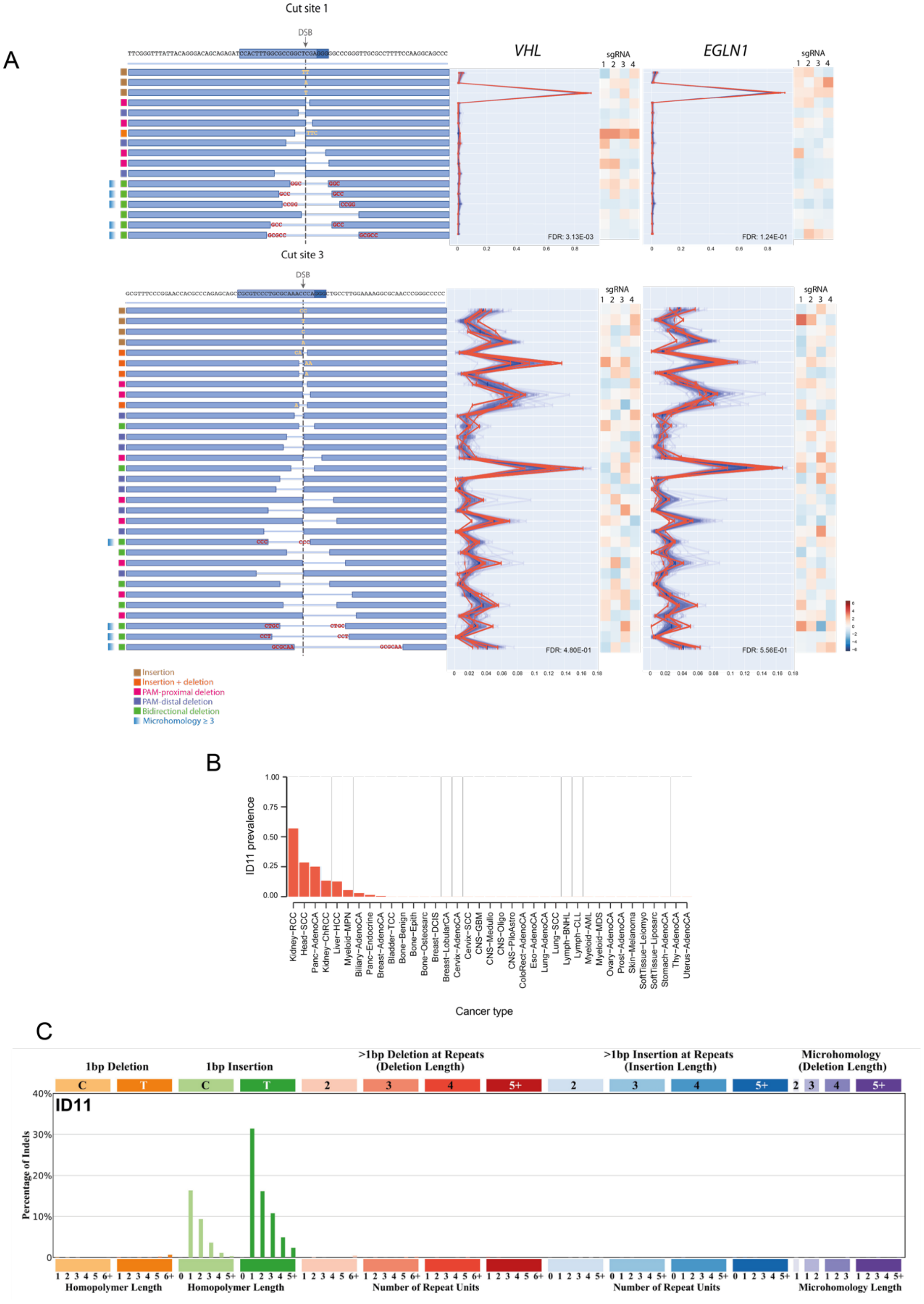
A
Cut site 1
DSB
VHL
EGLN1
sgRNA
1 2 3 4
FDR: 3.13E-03
FDR: 1.24E-01
Cut site 3
FDR: 4.80E-01
FDR: 5.56E-01
Insertion
Insertion + deletion
PAM-proximal deletion
PAM-distal deletion
Bidirectional deletion
Microhomology ≥ 3
B
ID11 prevalence
Cancer type
Kidney−RCC
Head−SCC
Panc−AdenoCA
Kidney−ChRCC
Liver−HCC
Myeloid−MPN
Biliary−AdenoCA
Panc−Endocrine
Breast−AdenoCA
Bladder−TCC
Bone−Benign
Bone−Epith
Bone−Osteosarc
Breast−DCIS
Breast−LobularCA
Cervix−AdenoCA
Cervix−SCC
CNS−GBM
CNS−Medullo
CNS−Oligo
CNS−PiloAstro
ColoRect−AdenoCA
Eso−AdenoCA
Lung−AdenoCA
Lung−SCC
Lymph−BNHL
Lymph−CLL
Myeloid−AML
Myeloid−MDS
Ovary−AdenoCA
Prost−AdenoCA
Skin−Melanoma
SoftTissue−Leiomyo
SoftTissue−Liposarc
Stomach−AdenoCA
Thy−AdenoCA
Uterus−AdenoCA
C
1bp Deletion
1bp Insertion
>1bp Deletion at Repeats (Deletion Length)
>1bp Insertion at Repeats (Insertion Length)
Microhomology (Deletion Length)
ID11
Percentage of Indels
Homopolymer Length
Homopolymer Length
Number of Repeat Units
Number of Repeat Units
Microhomology Length

**Fig. S7. *VHL* and *EGLN1* screen data and ID11 cancer signature.**

**(A)** Data from screen for *VHL* and *EGLN1* in TKOv3 Cut 1 and Cut 3, as in Fig. 1C.

**(B)** Relative frequency (i.e., prevalence) of tumour samples with detectable ID11 across 37 different tumour types and 2,608 tumour samples included in PCAWG (Pan Cancer Analysis of Whole Genomes). Tumour types are indicated in the x axis and the relative frequency is indicated in the y axis.

**(C)** Cancer mutational signature ID11 representation showing relative abundance of each type of indel depending on the sequence context. Image taken from COSMIC webpage (https://cancer.sanger.ac.uk/signatures/id/).

| | Polyclonal KO (%) | |
|---|---|---|
| **Gene** | **gRNA1** | **gRNA2** |
| *XLF* | 48 | 84 |
| *PAXX* | 38 | 40 |
| *POLL* | 65 | 84 |
| *NDST1* | 93 | 93 |
| *GET3* | 80 | 77 |
| *ATP6V1B2* | 84 | 86 |
| *HLTF* | N/A | 91 |
| *PCNA* | N/A | 85 |
| *VHL* | N/A | 94 |
| *HIF1a* | N/A | 95 |
| *HIF2a* | N/A | 96 |

**Table S3. Percentage of knockout, estimated by TIDE analysis, achieved by the indicated gene-targeting gRNAs in a polyclonal cell population.**

| | **Allele 1** | **Allele 2** |
|---|---|---|
| ***HLTF$^{-/-}$* Cas9 pLCKO2-NT** | | |
| clon #9 | +1nt at position 218 | +1nt at position 218 |
| clon #11 | +1nt at position 218 | -2nt at position 217 |
| clon #12 | +2nt at position 218 | +2nt at position 218 |
| ***HLTF$^{-/-}$* Cas9** | | |
| clon #7 | +4nt at position 218 | -2nt at position 216 |
| clon #9 | +1nt at position 218 | -2nt at position 216 |
| clon #10 | +1nt at position 218 | +1nt at position 218 |
| ***POLL$^{-/-}$* Cas9 pLCKO2-NT** | | |
| clon #5 | +1nt at position 229 | -7nt at position 222 |
| clon #10 | +1nt at position 229 | -1ntnt at position 228 |
| ***XLF$^{-/-}$* Cas9 pLCKO2-NT** | | |
| clon #3 | -14nt at position 26 | -14nt at position 26 |
| clon #21 | -1nt at position 25 | +1nt at position 26 |
| ***PAXX$^{-/-}$* Cas9 pLCKO2-NT** | | |
| clon #2 | +1nt at position 72 | +1nt at position 72 |
| clon #3 | +1nt at position 72 | +1nt at position 72 |
| ***GET3$^{-/-}$* Cas9 pLCKO2-NT** | | |
| clon #3 | -5nt followed by +1nt insertion at position 186 | -5nt followed by +1nt insertion at position 186 |
| clon #7 | -13nt at position 180 | -78nt at position 117 |
| ***POLQ$^{-/-}$* Cas9 pLCKO2-NT** | | |
| clon #3 | +5nt at position 137 | -11nt at position 131 |
| clon #7 | -4nt at position 133 | -19nt at position 130 |

**Table S5. Genotype of RPE-1 knockout clones generated in this work. Detected indels are indicated according to their position in the gene CDS sequence.**

| Guide RNA name | Guide RNA sequence (without PAM) |
|---|---|
| **Knockout generation** | |
| PAXX_1 | 5’ CTTCGCAGTAGCACACGAAG 3’ |
| PAXX_2 | 5’ GCTTCGTGTGCTACTGCGAA 3’ |
| XLF_2 | 5’ ACGCCCATGGCTGCATCAAC 3’ |
| XLF_3 | 5’ TGAACAGGTGGACACTAGTG 3’ |
| ASNA1-1 | 5’ CAGAACACTCTCACGCCCCT 3’ |
| ASNA1-2 | 5’ GCATCTGAGATGTTGTGTGC 3’ |
| ATP6V1B2_1 | 5’ GATGCGGGGGATTGTCAACG 3’ |
| ATP6V1B2_2 | 5’ GCGATGCGGGGGATTGTCAA 3’ |
| NDST1_1 | 5’ CGGCCTACTACCTATATGGC 3’ |
| NDST1_2 | 5’ ATCTCGGCCTACTACCTATA 3’ |
| HLTF_2 | 5’ GTTGGACTACGCTATTACAC 3’ |
| POLL_1 | 5’ GTGAGTGACACCTGGGCCCT 3’ |
| POLL_2 | 5’ AGGAGGCTGGTGGATGTAGC 3’ |
| PCNA_1 | 5’ GACCAGGCGCGCCTCGAACA 3’ |
| PCNA_2 | 5’ GCTCTGCAGGTTTACACCGC 3’ |
| VHL_1 | 5’ CATACGGGCAGCACGACGCG 3’ |
| VHL_2 | 5’ GAACTCGCGCGAGCCCTCCC 3’ |
| HIF1A_1 | 5’ AACCATAACAAAACCATCCA 3’ |
| HIF1A_2 | 5’ TTCTCCACTTAGGAGTAGCT 3’ |
| POLQ_1 | 5’ CGGCCTTCAGGCAGCGACCA 3’ |
| **Repair profile analysis** | |
| AAVS1 site 1 | 5’ CCTCTAAGGTTTGCTTACGA 3’ |
| AAVS1 site 2 | 5’ CTCCCTCCCAGGATCCTCTC 3’ |
| AAVS1 site 3 | 5’ TGGGGGTGTGTCACCAGATA 3’ |
| AAVS1 site 4 | 5’ CATCCTAACGTCCTTTTCCC 3’ |
| TKOv3 site 1 | 5’ CCACTTTGGCGCCGGCTCGA 3’ |
| TKOv3 site 2 | 5’ CCCGGGTTGCGCCTTTTCCA 3’ |
| TKOv3 site 3 | 5’ CGCGTCCCTGCGCAAACCCA 3’ |
| TKOv3-Non targeting | 5’ GTGAACCGCATCGAGCTGAA 3’ |
| CloneTracker site 1 | 5’ CTTTTCTTTTAAATTGCTCG 3’ |
| CloneTracker site 2 | 5’ CTCGAGCAATTTAAAAGAAA 3’ |
| Multi-site | 5’ CCTGTAGTCCCAGCTACTCT 3’ |

**Table S7. List and sequence of guide RNAs used in this work.**

| Gene | Sequence (5’-3’) |
|---|---|
| NDST1_Fwd | GCTGGGAGGAAGGACTGG |
| NDST1_Rev | GCCCTTGTCAGTGAGCGT |
| NDST1_Seq | TCTCGTGTCTCCTCCTGTGTGG |
| ASNA1_E2_Fwd | CCACCATAACCCATACTCTCCT |
| ASNA1_E2_Rev | CCCTACCCTGGCTCACTAAG |
| ASNA1_E2_Seq | GGATATAGAGTGGATAGGACACGG |
| ATP6V1B2_Fwd | CTCTCTTTCGGCTGGCTTTC |
| ATP6V1B2_Rev | ACAAGTGGGTCAGGGTCG |
| ATP6V1B2_Seq | AAGCAGGTCTGGCCCCAGCG |
| XLF_E1_Fwd | GCTGGTTGTATACTGGGAGAACG |
| XLF_Rev | ACTGGAATTGTCTCAACCCGA |
| XLF_Seq | CAGACTGGTTTGGGTGGTGC |
| PAXX_Fwd | AAGTCGTTCTTCCGCGCAG |
| PAXX_V2.1_Rev | CCCCAGTGAAGGTATGAGGT |
| PAXX_V2_Seq | GTTACCCACGAGGGCCGCCAG |
| POLL_E3_Fwd | AGGAAAGCTGTCTGGATGGA |
| POLL_E3_Rev | AGGGTCTTGCTCTGTCACC |
| POLL_E3_Seq | AAGTTATGTCGTGGCCAGGGG |
| HLTF _P2_Fwd | GTGAACTCTGACTAGGCTTTTCA |
| HLTF _P2_Rev | CCAAACATAACGCACTGAGGC |
| VHL_E1_Fwd | GTGGTCTGGATCGCGGAG |
| VHL_E1_Rev | CCCCGTCTGCAAAATGGAC |
| VHL_E1_Seq | GGAGAACTGGGACGAGGCC |
| HIF1a_Fwd | TGGCCTGCACTTTTCTGTGTT |
| HIF1a_Rev | CTCATGGTCACATGGATGAGTAAA |
| HIF2a_Fwd | ATGCAAGAGGAGGACTTTGGT |
| HIF2a_Rev | TGGGTGAAAGTTCTGTAGTGGG |
| POLQ_Fwd | AGACTGGCGGGAAGATGTC |
| POLQ_Rev | CAGTCACAGAGAAGGGGAGT |
| POLQ_Seq | ACTGTGGCTTGCCCTGATCGG |
| ASNA1qpcr_Fwd | ACAGACCCAGCACACAACAT |
| ASNA1qpcr_Rev | CAGCATGTTGTCCTCCTCGA |

**Table S8. Primers used for gene knockout validation.**

| Target_site | Sequence (5'-3') |
|---|---|
| AAVS1_1_Fwd | GCAGCACCAGGATCAGTGAA |
| AAVS1_1_Rev | ATTCCCAGGGCCGGTTAATG |
| AAVS1_2_Fwd | TGACGCACGGAGGAACAATA |
| AAVS1_2_Rev | GCCACTAGGGACAGGATTGG |
| pLCKO2_Fwd | CAGTGCAGGGGAAAGAATAGTAGACA |
| pLCKO2_Rev | GGAACAGGGCCCACACTACC |
| pLCKO2_cut2_seq | CAAGGAACCTTCCCGACTTAG |
| GREB1_3_Fwd | CTTGCTTCTGAAGGGCAGAG |
| GREB1_3_Rev | GACCCAGTTGCCACACTTTT |
| cutting efficeincy_qPCR_Fwd | TCGGGTTTATTACAGGGACAGC |
| cutting efficeincy_qPCR_Rev | CGTGAAGAATGTGCGAGACC |

**Table S9. Primers used for repair analysis.**
